# Revisiting the evolution and taxonomy of Clostridia, a phylogenomic update

**DOI:** 10.1101/546341

**Authors:** Pablo Cruz-Morales, Camila A. Orellana, George Moutafis, Glenn Moonen, Gonzalo Rincon, Lars K Nielsen, Esteban Marcellin

## Abstract

*Clostridium* is a large genus of obligate anaerobes belonging to the *Firmicutes* phylum of bacteria, most of which have a Gram-positive cell wall structure. The genus includes significant human and animal pathogens, causative of potentially deadly diseases such as tetanus and botulism. Despite their relevance and many studies suggesting that they are not a monophyletic group, the taxonomy of the group has largely been neglected. Currently, species belonging to the genus are placed in the unnatural order defined as *Clostridiales*, which includes the class *Clostridia*. Here we used genomic data from 779 strains to study the taxonomy and evolution of the group. This analysis allowed us to; (i) confirm that the group is composed of more than one genus (ii), detect major differences between pathogens classified as a single species within the group of authentic *Clostridium* spp. (*sensu stricto*), identify inconsistencies between taxonomy and toxin evolution that reflect on the pervasive misclassification of strains and, (iv) identify differential traits within central metabolism of members of what has been defined earlier and confirmed by us as cluster I. Our analysis shows that the current taxonomic classification of *Clostridium* species hinders the prediction of functions and traits, suggests a new classification for this fascinating class of bacteria and highlights the importance of phylogenomics for taxonomic studies.

## Introduction

Clostridia are an important genus of Gram-positive, often anaerobic, rod shaped, spore-forming bacteria. The group includes important human and animal pathogens such as *C. botulinum, C. tetani, and C. difficile* as well as industrially relevant microorganisms such as *C. acetobutylicum*. The importance of the genus is reflected by the more than 42,000 entries in the Pubmed database, and about 1,700 genome sequences from this group deposited in the GenBank database.

Early molecular analyses in the 1970s demonstrated considerable diversity and ambiguities among the genus (Johnson and Francis, 1975), In fact, this early classification of the genus *Clostridium* does not respect the identity thresholds established for 16s rRNA (Rossi-Tamisier et al, 2015), a widely used taxonomic marker. In consequence, this classification has been revisited several times (Collins *et al.,* 1994; Yutin and Galperin, 2013; Lawson, 2016a). Currently, it is well known that there are at least three *C. botulinum* lineages and that *C. difficile* belongs to a distantly related genus leading to the recent reclassification of *C. difficile* as a C*lostridioides difficile* (Lawson *et al.*, 2016b). However, the taxonomic relationships and evolutionary dynamics of the species associated with the genus remain largely neglected.

The recent availability of sequenced genomes provides a new opportunity to revisit its taxonomy in the genomic era as opposed to taxonomic classification based on 16S rRNA sequencing. Such an opportunity enables a comprehensive taxonomic and evolutionary analysis to confirm that they are not a monophyletic group and there is a need to redefine the group taxonomically.

In this work, we have compiled the genomes classified as “*Clostridium”* and “*Clostridioides*” in the GenBank database (Benson *et al.*, 2006) to identify a set of conserved genes to be used to define taxonomy. Once the classification was established, we focused on what has been called “cluster I” species (*sensu stricto*) (Lawson *et al.*, 2016a) to identify differences between the core/pan genomes of cluster I strains and to reveal general evolutionary trends and specific traits linked to adaptation to different lifestyles.

## Results and discussion

### 1. General Taxonomy

We first retrieved more than 1,700 genomes and draft genomes deposited as “*Clostridium”* and “*Clostridioides*” from the GenBank as of July 2017. The dataset was filtered by removing low quality genomes (genomes with more than 400 contigs) and by eliminating redundancy at the strain level. This filtering resulted in a subset of 779 genomes (Supplementary Table S1) used hereafter. We used the taxonomic definition of clostridial “clusters” as reference and annotated those strains with a species name accordingly (Rainey *et al*., 2006; Bowman *et al*., 2010; Jung *et al*., 2010, Liou *et al*., 2005; Sakuma *et al*., 2006; Shiratori *et al*., 2009; Slobodkina *et al*., 2008; Tamburini *et al*., 2001; Warren *et al*., 2006).

From this subset of genomes, we identified 27 conserved protein sequences (Supplementary Table S2). These proteins do not represent the complete core genome of clostridia, instead, these proteins are a set of taxonomic markers that can confidently be used for the construction of a clostridial species tree (Figure 1). This tree defined 7 major clades that were consistent with the previous established clostridial “clusters” classification (Rainey *et al*., 2006). Accordingly, clusters III, IV, Xia, XIVa, XIVb, and XVI were distantly related to Cluster I (*sensu stricto*), which contains 370 strains including most toxin-producer pathogens and industrially relevant strains, but clearly excludes *difficile* species. Based on this analysis we defined the members of cluster I as the authentic members of the *Clostridium* genus. Furthermore, since cluster Xia has already been reclassified as *Clostridioides* sp. we propose that the remaining 5 groups represent distinct new genera.

**Figure 1.**
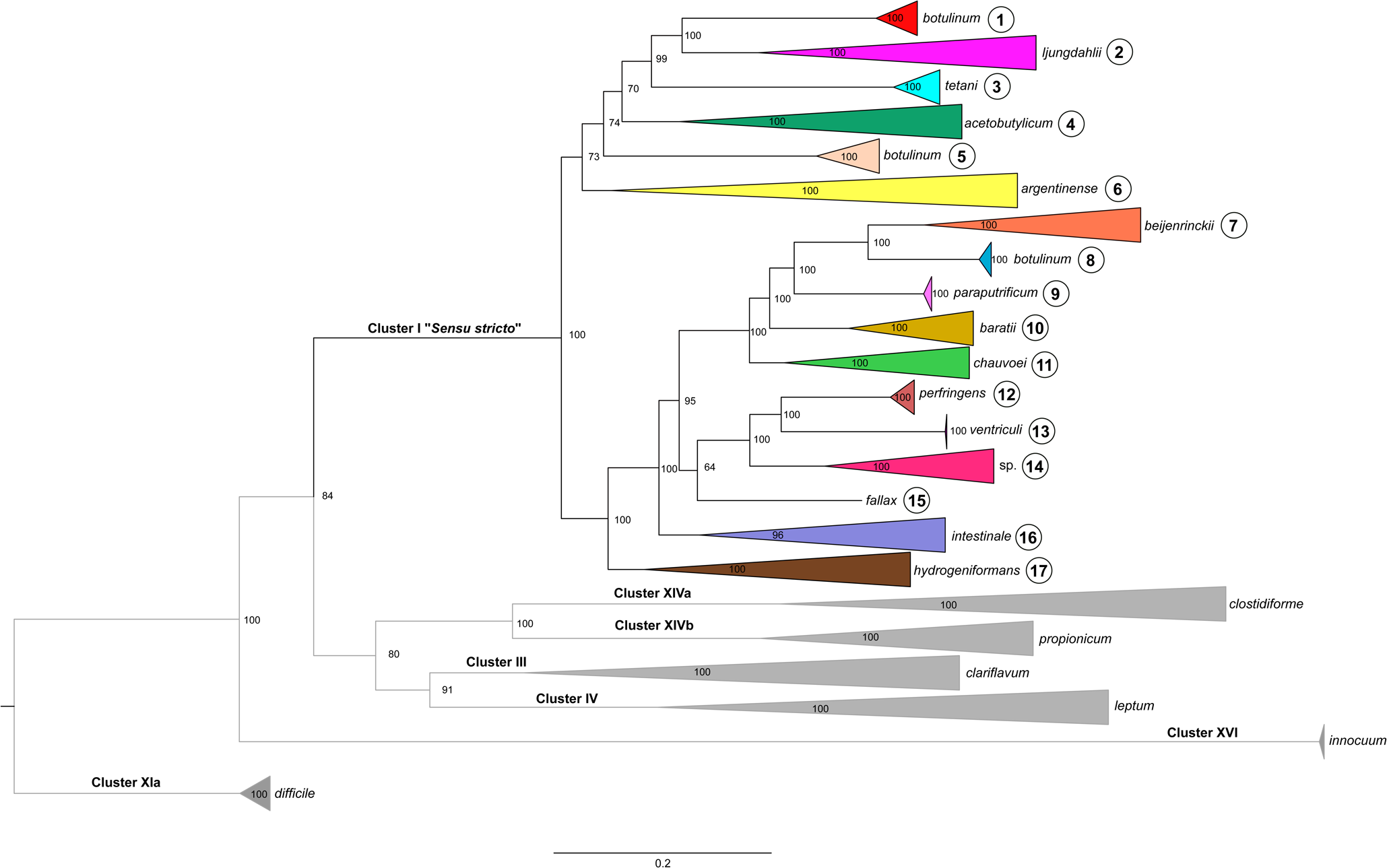
Phylogenetic reconstruction of *Clostridium* species. This phylogeny was constructed using 27 markers conserved across 779 genomes (Supplementary Table S1) deposited in the GenBank database and taxonomically defined as *Clostridium*. The main clades outside and within (370 taxa) the *sensu stricto* group (real clostridia) have been collapsed and defined as 17 taxonomic subgroups (Table 1). Branch support is shown at each node. Uncollapsed clades for subgroups 1-17 are shown in Supplementary Figures S1-S14.

Cluster I was further divided into 17 subgroups (Table 1) using the species tree presented in Figure 1 Our analysis also showed that strains named *C. botulinum* are found in subgroups 1, 5 and 8 (Table 1). These clades include *C. botulinum* strains defined by their toxin types as A / B / F (subgroup 1), C / D / CD (Subgroup 5) and E (subgroup 8). Clades 1 and 5 also include other species namely: *C. sporogenes* in Clade 1 and C. *haemolyticum*, and *C. novyi* in Clade 5.

**Table 1.** Subgroup Core genome analysis.

| Subgroup | Number of strains | Species | Proteins in Subgrocore | Core of cores |  |  |
| --- | --- | --- | --- | --- | --- | --- |
|  |  |  |  | Proteins in core | Accessory proteins | Unique proteins |
| 1 | 106 | *botulinum* A, B, F, *sporogenes* | 890 | 212 | 673 | 2 |
| 2 | 21 | *autoethanogenum*,<br>*carboxidivorans*, *coskatii*,<br>*drakei*, *kluyveri*,<br>*ljungdahlii*, *magnum*, *ragdalei*,<br>*scatologenes*, *tyrobutyricum* | 1071 | 212 | 755 | 99 |
| 3 | 10 | *tetani* | 2214 | 212 | 1115 | 877 |
| 4 | 16 | *acetobutylicum*, *akagii*, *arbusti*,<br>*aurantibutyricum*, *felsineum*,<br>*pasteurianum*, *roseum* | 1063 | 212 | 752 | 96 |
| 5 | 42 | *botulinum* C & D, *haemolyticum*,<br>*novyi* | 1367 | 212 | 824 | 324 |
| 6 | 11 | *argentinese*, *collagenovorans*,<br>*estertheticum*, *proteolyticum*,<br>*senegalense*, *sulfidigenes*,<br>*tepidiprofundii*, *tunisiense* | 462 | 212 | 245 | 4 |
| 7 | 48 | *beijerinckii*, *butyricum*,<br>*puniceum*, *saccharobutylicum*,<br>*saccharoperbutylacetonicum* | 1319 | 212 | 925 | 178 |
| 8 | 20 | *botulinum* E | 994 | 212 | 769 | 10 |
| 9 | 4 | *paraputrificum* | 2704 | 212 | 1452 | 1021 |
| 10 | 8 | *baratii*, *colicanis* | 1350 | 212 | 1037 | 94 |
| 11 | 15 | *chauvoei*, *disporicum*,<br>*sartagoforme*, *saudiense*,<br>*septicum* | 1028 | 212 | 779 | 32 |
| 12 | 55 | *perfringens* | 2044 | 212 | 1276 | 547 |
| 13 | 3 | *ventriculi* | 2040 | 212 | 1091 | 732 |
| 14 | 2 | Spp. | 1505 | 212 | 1043 | 242 |
| 15 | 1 | *fallax* | 2415 | 212 | 1336 | 854 |
| 16 | 3 | *cavendishii*, *intestinale* | 1500 | 212 | 1104 | 178 |
| 17 | 5 | *algidicarnis*, *cadaveris*,<br>*hydrogeniformans* | 992 | 212 | 719 | 58 |

Comparison of the overall synteny between *C. botulinum* strains from Clades 1, 5 and 8 showed divergence among them. High synteny could be observed between *C. botulinum* strains from Clade 1 and *C. sporogenes* as well as *C. botulinum* strains from Clade 5 and *C. novyi*, respectively (Supplementary Figure S15). The fact that *C. sporogenes, C. novyi* and *C. haemolyticum* species show little divergence with their respective *C. botulinum* relatives suggest that these strains are either artificially defined as distinct species or have just recently diverged. Together, these observations indicate that the strains defined as *Clostridium botulinum* should be split into three species found within groups 1, 5 and 8. *C. botulinum* strains in subgroups 1 and 5 may be called, *C. sporogenes* and *C. haemolyticum,* respectively, since these species have been previously defined, while strains within subgroup 8 may remain as members of the authentic *C. botulinum* species. However, as highlighted by Lawson (2016b), changing names of medically relevant organisms can cause great confusion in the healthcare community. As these three species produce botulinum neurotoxins, the change of name might be rejected under Rule 56a (5) of the International Code of Nomenclature of Prokaryotes (Parker, Tindall, & Garrity, 2015), which states that “names whose application is likely to lead to accidents endangering health or life or both” can be rejected.

As this analysis uses draft genomes to include as many genomes as possible, and only 27 proteins were conserved among these genomes, the analysis was repeated using a smaller subset of high quality genomes (179, N50>600 kbp) to validate our results. As such, a higher number of conserved proteins (79) were obtained and used (Supplementary Table S3). This new analysis (Supplementary Figure S16) showed that the taxonomic groups maintained the same distribution (tree topology) when using a dataset of 179 or 779 genomes and a matrix containing 79 or 27 protein sequences respectively. The same Clusters and Cluster I subgroups were observed (Supplementary Figure S16-S29), with the exception of clades that disappeared as they did not pass the stringent genome quality cut off (Cluster IV and XVI, and Subgroups 9, 13, 14 and 15 in Cluster I).

### 2. Toxin evolution

Pathogenic clostridia produce the highest number of life-threatening toxins of any genus. This includes enterotoxins that affect the gut, such as *C. difficile* toxins A and B, histotoxins that affect soft tissue such as *C. perfringens* and *C. septicum* alpha-toxins, and neurotoxins affecting nervous tissue such as tetanus (*C. tetani*) and botulinum (*C. botulinum*) toxins. Diseases range from gastroenteritis to abdominal disorders, colitis, muscle necrosis, soft tissue infections, tetanus and botulism amongst others (Hatheway, 1990). These toxin-encoding genes are often located on mobile genetic elements or in variable regions of the chromosome (Hatheway, 1990; Petit, Gibert, & Popoff, 1999; Skarin & Segerman, 2011), resulting in gene transfer between species. Here we analyzed different toxins evolution to compare taxonomy with phylogeny.

The botulinum neurotoxin (BotA) for example, represents the most poisonous biological protein known and has been used as a phenotypic and genotypic marker for taxonomic classification. In fact, *C. botulinum* strains are often classified as members of groups A, B, C, D, E and F, in direct relationship with the production of antigenically distinguishable variants of the neurotoxin. In this work, homologs of BotA were found exclusively amongst members of Cluster I, and were distributed amongst *C. botulinum, C. tetani, C. argentinense, C. baratii* and *C. butyricum* species. A phylogenetic reconstruction of these homologs (Figure 2A) showed little divergence except for three homologs; two on *botulinum* species, and one on *C. argentinense* that seem to be more divergent.

**Figure 2.**
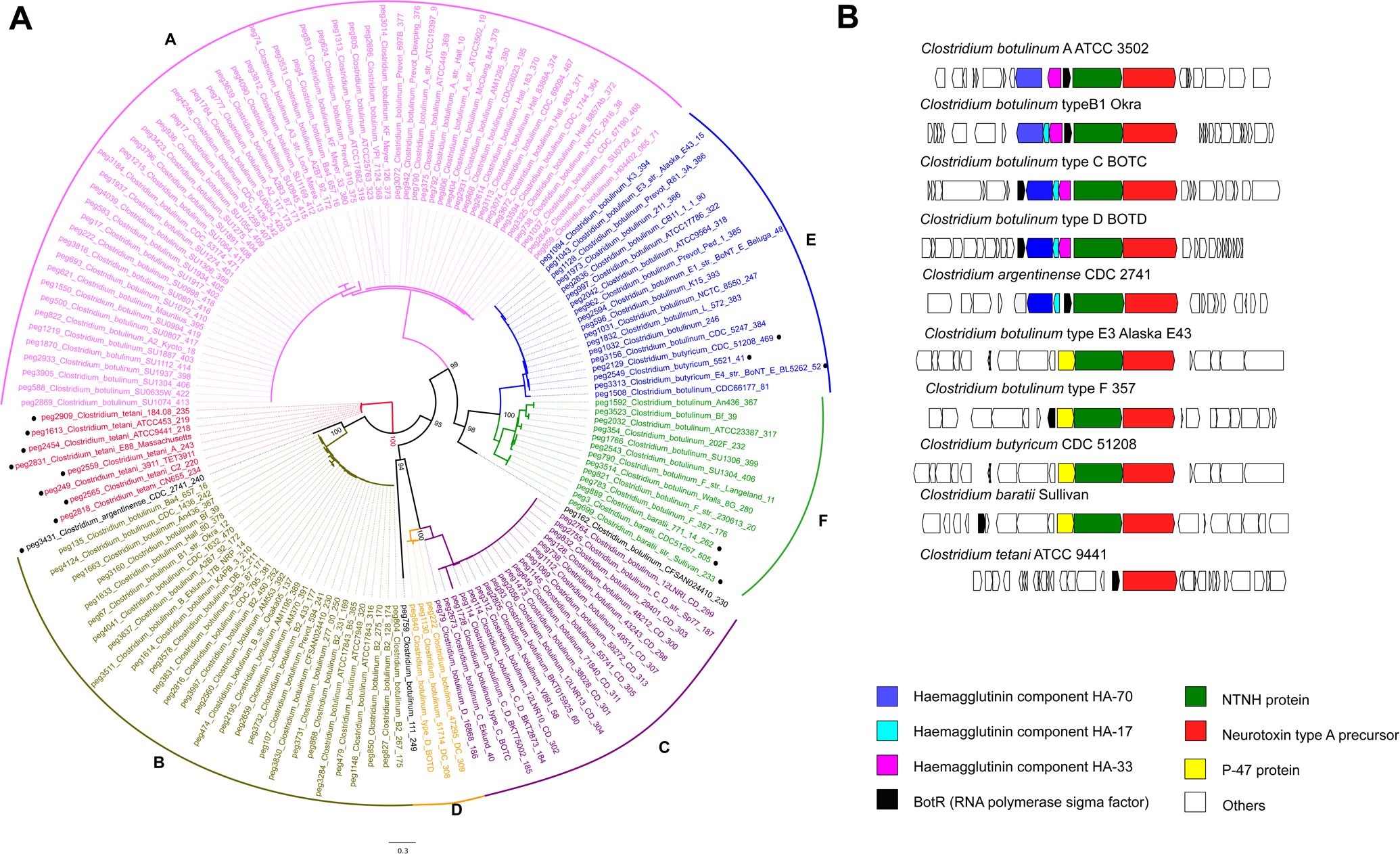
A. Phylogenetic reconstruction of BotA toxin proteins. Six A-F clades are consistent with previous reports. Including non-*botulinum* strains, *argentinense, tetani, butyricum* and *baratti* (marked with a dot). Three new sequences (in black) account for new unclassified toxin diversity. **B.** Genome context of BotA homologs found in Cluster I strains.

The topology of the BotA phylogeny agrees with previous definitions of the *C. botulinum* subgroups A, B, C, D, E, and F, with clades populated by strains with similar toxin types (i. e. Clade A has only *C. botulinum* A strains, etc.). However, toxin markers were not consistent with the species tree, for which *C. botulinum* toxins types A, B and F were in Clade I while grouping independently in the toxin tree.

A gene context analysis (Figure 2B) showed two major synteny groups {A, B, C, D} and {E, F}. The presence of toxin accessory proteins (Lam *et al*., 2017) was found to be the main difference between them, namely haemagglutinin coding genes in groups A, B, C, and D, and protein p47 in groups E and F. These observations are consistent with *C. botulinum* strains located in subgroups 1, 5 and 8 being distinct species that acquired the toxin genes by horizontal gene transfer hindering taxonomic classification.

The analysis of other important toxins also shows many horizontal gene transfer events of toxin genes between subgroups. *C. difficile* toxins A and B homologs were distributed amongst *C. difficile, C sordellii, C. acetobutylicum* and *C novyi* species (supplementary Figure S30). Homologs of *C. perfringens* alpha toxin were observed in *C. perfringens, C. novyi, C. botulinum* C and D, *C. baratii, C. hemolyticum, C. cavendishii, C. argentinense, C. sordellii* and *C. dakarense* species (supplementary Figure S31). Finally, *C. septicum* toxin alpha homologs were distributed amongst *C. septicum, C. novyi, C. haemolyticum* and *C. botulinum* C and D species (supplementary Figure S32). A summary of these findings can be found on Supplementary Table S4. Interestingly, *C. botulinum* C and D (subgroup 5) also have *C. perfringens* and *C. septicum* alpha toxins orthologs, while *C. botulinum* A, B, E and F do not. According with these observations, we suggest that toxin production should not be used to define taxonomic groups, as it uncouples taxonomy from phylogeny.

### 3. Core Genome analysis

Once the taxonomic framework was established, we used it to study the evolutionary dynamics and to identify general differences among the subgroups within Cluster I. For this purpose, we calculated core/pangenomes for each subgroup having more than 10 genomes (Table 1). This analysis (Figure 3) showed that subgroups 3 (*C. tetani),* 5 (*C. botulinum* toxin group C and D, *C. haemolyticum*, and *C. novyi*), 8 (*C. botulinum* toxin group E) and 12 (*C. perfringens*) have almost closed pangenomes, implying loss of genetic diversity. This observation is consistent with the evolutionary dynamic observed in pathogenic species by other authors attributed to “Specialist” species (Georgiades and Raoult, 2011).

**Figure 3.**
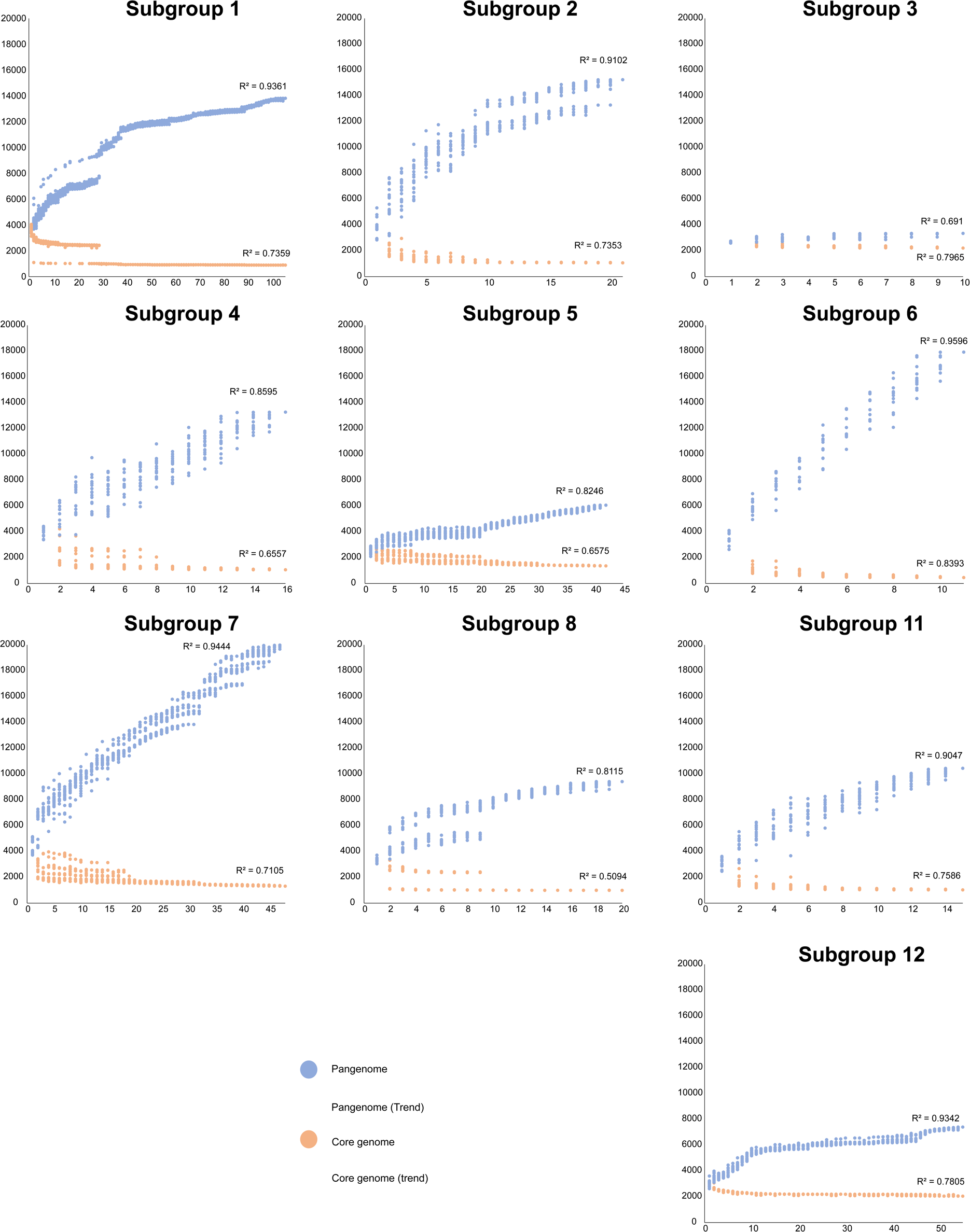
Pangenome analysis of selected subgroups. The **Y** axis shows the number of gene families and the **X** axis shows the number of genomes analysed. The number of conserved genes was calculated by randomly adding genomes, with 20 replicates (if n>20) or the same numbers as genomes (if n<20). This analysis shows large differences in the genetic diversity of the subgroups, with less diversity and almost closed pangenomes in pathogenic sub groups.

In contrast, the remaining lineages showed open pangenomes. Subgroup 1, which includes important pathogens such as *C. botulinum* toxin groups A, B and F, and the closely related C. *sporogenes* strains, have open pangenomes, implying larger genetic diversity and probably more recent adaptation to a pathogenic life-style. These observations further emphasize the presence of three distinct lineages among *C. botulinum* strains that may be re-classified as distinct species.

After establishing general differences between the evolutionary dynamics of the subgroups, we took a closer look at differences at the functional level. For this purpose, we extracted amino acid sequences of the core genes of each subgroup and identified conserved functions among them (i. e. a core of cores) and functions that are distinctive of each subgroup (Table 1). This analysis revealed that Cluster I has a core of 212 genes. As expected, many of these conserved genes are associated with housekeeping functions such as nucleotide biosynthesis, replication and repair (Supplementary Figure S33). Unique genes were abundantly classified as members of carbohydrate metabolism and for membrane transport. Interestingly, the largest number of accessory functions were related to amino acid metabolism, implying that multiple genes for this category are conserved at the subgroup level only. This observation is illustrated by the example described in the following section.

### 4. Divergence of the shikimate pathway in pathogenic clostridia

To investigate adaptive traits that could define differences within each sub group, we mined the pangenomes for functions that were uniquely found in each group. From a taxonomic point of view, unique genomic traits are important as they can be used for the development of genetic markers and to identify distinctive phenotypes that can be used for classification. The rationale for searching unique traits within subgroups was that the use of such a large genomic database would enable, for the first time, to find unique functions conserved in all members of a subgroup but absent in other subgroups, thereby enabling to dissect for subgroup specific adaptations. This was the case for the essential enzyme 3-phosphoshikimate 1-carboxyvinyltransferase (AroA), which was found in the pangenome of Cluster I. AroA is part of the shikimate pathway and is essential for the biosynthesis of aromatic amino acids phenylalanine, tyrosine and tryptophan. This seemed unusual given that all subgroup cores include AroA.

We reasoned that the presence of AroA among the pangenome may be due to; (i) divergence among AroA orthologs beyond the cutoff for orthology defined in our pangenome strategy leading to fragmentation of the gene family, or (ii) duplication events in certain subgroups and divergence, which have been previously linked to adaptive evolution in bacteria (Schniete *et al*., 2018). To explore this idea, we searched for homologs of the AroA enzyme in all the strains from Cluster I and found a single ortholog conserved in most strains. Thus, we assumed that AroA has divergently evolved within the Cluster I species.

Phylogenetic reconstruction of AroA (Figure 4A) confirmed the presence of two largely divergent AroA clades, one including homologs from strains in most subgroups and the other including subgroups 1,3 and 11. Interestingly, most strains in this clade can colonise human hosts and are toxin producing pathogens, except for *C. sporogenes*. Inspection of the genome context of representative AroA homologs from different subgroups (Figure 4B) revealed that despite sequence divergence, AroA homologs from *C. tetani, C. botulinum* toxin groups A, B and F, and *C. sporogenes* are located within a gene neighborhood that includes enzymes from the shikimate pathway. Thus, the gene context topology indicates that the function of these enzymes is linked to the production of aromatic amino acids. However, the divergent AroA homologs were found associated with the pyrimidine-associated regulator *pyrR* and a uracil permease. Such genomic organisation, suggests a link between aromatic amino acid biosynthesis and pyrimidine utilisation. A recent in-depth molecular characterisation of the *C. tetani* toxin production fermentation showed a potential link between extracellular uracil concentration and toxin production (Licona-Cassani *et al.*, 2016). However, this link is yet to be fully understood.

**Figure 4.**
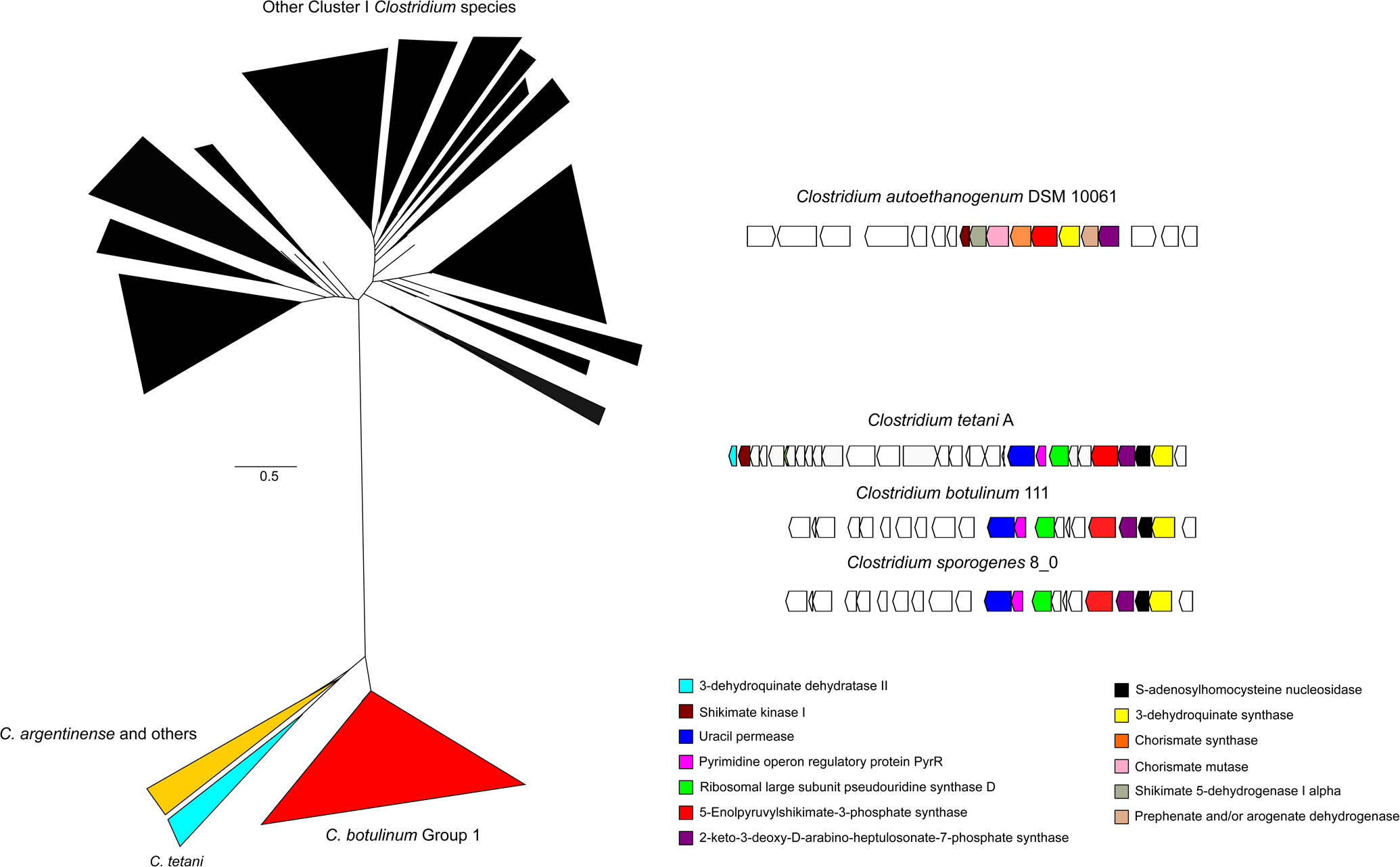
Phylogenomic analysis of AroA in Clostridium Group I. **A** The phylogeny shows that AroA, has significantly diverged in all members of the subgroup 3 (*C. tetani;* clear blue), subgroup 6 (*C. argentinense*; yellow) and subgroup 1 (*C. sporogenes* - *C. botulinum* B; red) from the rest of the subgroups in Cluster I (black). The full tree is provided as Supplementary Tree 2. **B. Genome context of AroA homologs.** *C. autoethanogenum* is shown as a typical Group I AroA genome context while divergent homologs show a genome context that includes enzymes from pyrimidine metabolism.

Studies have also shown that *C. sporogenes*, a soil bacterium rarely pathogenic for humans (Inkster et al, 2011) although it may be found in the gut, and *C. botulinum* (Cluster I), a toxin producing pathogen, copiously produce tryptophan, phenylalanine and tyrosine. It has been suggested that secretion of these amino acids and intermediates of its degradation may influence intestinal permeability and systemic immunity of the host (Dodd *et al*., 2018). We speculate that the divergence in AroA may be related to the evolution of new metabolic interactions that do not affect the enzymatic activity of AroA, but rather its regulation in clostridial species that are able to colonise hosts. Given the presence of this trait in pathogen and commensal strains, we reasoned that this trait likely evolved prior to the acquisition of toxin genes. Following the same thought process, we suggest that *C. botulinum* subgroup 1 toxin groups A, B and F, have only recently evolved into pathogenic organisms. By selecting amino acid biosynthesis to illustrate the use of the new classification, we show here that a correlation between traits, function gain and loss cannot be extracted from the current taxonomic classification of *Clostridium* species, and they remain unchanged. Through this effort, we hope that our work serves to inspire the research community to study the evolution of clostridia at the genome-scale level and suggest a new classification for this fascinating class of bacteria.

## Conclusions

Here we present an inclusive framework for phylogenomic analysis aimed at providing an updated view of the *Clostridium* genus. Our work shows that the current definition of clostridia encompasses a large and diverse group of species that is inconsistent with its definition as a genus. Instead, the group includes multiple genera. Furthermore, within group I, arguably the authentic *Clostridium* genus, further taxonomic inconsistencies exist due to the use of BotA for taxonomic classification as a taxonomic marker. This has previously been observed by others (Yutin and Galperin, 2013; Lawson 2016a; Lawson, 2016b; Weigand *et al*., 2015; Udaondo *et al*., 2017) but to the best of our knowledge, clostridial taxonomy and evolution has not been revisited using the opportunity offered by next generation sequencing for phylogenomic reclassification until now. Given the pervasiveness of the misclassification in clostridial species, we wonder whether the current system of classification should be kept, or if it should be revisited and simplified using genomic data. The recent explosion of available annotated genomes offers an unprecedented opportunity to answer intriguing questions surrounding pathogenic clostridial evolution. For example, the incredible diversity and the number of toxins produced by some strains is yet to be fully understood. So is the astonishing potency of some of the toxins produced by these pathogens, which must confer an evolutionary advantage that remains to be elucidated.

## Materials and methods

All genomes were downloaded from the NCBI FTP site, filtered by the number of contigs, (cutoff <= 400) resulting in 779 genomes which were annotated in RAST (Aziz *et al*., 2008). The conserved proteins present in the selected genomes were identified using BPGA v1.3 (Chaudhari *et al*., 2016) with an identity cutoff of 0.4 for clustering of groups of orthologs using Usearch (Edgar, 2010).

The resulting 27 groups of orthologs were aligned using Muscle v3.8 (Edgar *et al*., 2004) and the alignments were manually curated and concatenated using SeaView v4 (Gouy *et al*., 2010). The final amino acid matrix included 12,836 amino acids. The best amino acid substitution model for each of the 27 partitions (Supplementary Table S1) was selected using the ModelFinder tool implemented in IQ-tree (Kalyaanamoorthy *et al*., 2017) and the phylogeny was constructed using IQ-tree (Nguyen *et al*., 2015), using the partitioned models with 10,000 bootstrap replicates.

Pangenome analysis of Cluster I subgroups was performed using BPGA following the same approach described above. Homologs of BotA and AroA were mined and retrieved from the database using BlastP (Altschul *et al*., 1990) with an e-value cutoff of 1E-9 and bit score of 200. Phylogenetic trees for clostridial toxins and AroA were obtained using the same approach. Synteny analysis was performed using CORASON-BGC (Cruz-Morales *et al*., 2017) with an e-value cutoff of 1E-9 and a bit score of 200.

The full non-collapsed aroA and species trees are available as Supplementary Tree 1 deposited at TreeBASE (Vos et al, 2012): http://purl.org/phylo/treebase/phylows/study/TB2:S23279

## Acknowledgments

PCM thanks the Australian Department of Education and Training for the Endeavour Fellowship. The authors thank Robin Palfreyman for technical support, Cuauhtemoc Licona-Cassani and Nicolas Zaragoza for helpful discussions and the anonymous reviewers for constructive criticism. This research was funded through an Australian Research Council Linkage Grant LP150100087 with Zoetis as the Industrial Partner. Elements of this research utilised support provided by the Queensland Node of Metabolomics Australia, an initiative of the Australian Government being conducted as part of the NCRIS National Research Infrastructure for Australia.

## References

1. Altschul SF, Gish W, Miller W, Myers EW, Lipman DJ. 1990. Basic local alignment search tool. J Mol Biol. 215(3):403–10.

2. Benson DA, Karsch-Mizrachi I, Lipman DJ, Ostell J, Wheeler DL. 2006. GenBank. Nucleic Acids Res. 34 (Database issue):D16–20.

3. Bowman KS, Dupré RE, Rainey FA, Moe WM. 2010. *Clostridium hydrogeniformans* sp. nov. and *Clostridium cavendishii* sp. nov., hydrogen-producing bacteria from chlorinated solvent-contaminated groundwater. Int J Syst Evol Microbiol. 60(Pt 2):358–63.

4. Chaudhari NM, Gupta VK, Dutta C. 2016. BPGA-an ultra-fast pan-genome analysis pipeline. Sci Rep. 13;6:24373.

5. Collins MD, Lawson PA, Willems A, Cordoba JJ, Fernandez-Garayzabal J, Garcia P, Cai J, Hippe H, Farrow JA. 1994. The phylogeny of the genus *Clostridium*: proposal of five new genera and eleven new species combinations. Int J Syst Bacteriol. 44(4):812–26.

6. Cruz-Morales P, Ramos-Aboites HE, Licona-Cassani C, Selem-Mójica N, Mejía-Ponce PM, Souza-Saldívar V, Barona-Gómez F. 2017. Actinobacteria phylogenomics, selective isolation from an iron oligotrophic environment and siderophore functional characterization, unveil new desferrioxamine traits. FEMS Microbiol Ecol. 93(9). fix086

7. Dodd D, Spitzer MH, Van Treuren W, Merrill BD, Hryckowian AJ, Higginbottom SK, Le A, Cowan TM, Nolan GP, Fischbach MA, Sonnenburg JL. 2017. A gut bacterial pathway metabolizes aromatic amino acids into nine circulating metabolites. Nature. 551(7682):648–652.

8. Georgiades K, Raoult D. 2011. Defining pathogenic bacterial species in the genomic era. Front Microbiol. 1:151.

9. Gouy M, Guindon S, Gascuel O. 2010. SeaView version 4: A multiplatform graphical user interface for sequence alignment and phylogenetic tree building. Mol Biol Evol. 27(2):221–4.

10. Hatheway, CL, 1990. Toxigenic clostridia. Clin Microbiol Rev. 3(1): 66–98.

11. Inkster T, Cordina C, Siegmeth A. 2011. Septic arthritis following anterior cruciate ligament reconstruction secondary to *Clostridium sporogenes*; a rare clinical pathogen. J Clin Pathol. 64(9):820–1.

12. Johnson JL, Francis BS. 1975. Taxonomy of the Clostridia: ribosomal ribonucleic acid homologies among the species. J Gen Microbiol. 88(2):229–44.

13. Kalyaanamoorthy, B.Q. Minh, T.K.F. Wong, A. von Haeseler, L.S. Jermiin. 2017. ModelFinder: Fast model selection for accurate phylogenetic estimates. Nat. Methods, 14:587–589.

14. Jung MY, Park IS, Kim W, Kim HL, Paek WK, Chang YH. 2010. *Clostridium arbusti* sp.nov., an anaerobic bacterium isolated from pear orchard soil. Int J Syst Evol Microbiol. 60(Pt 9):2231–5.

15. Lam KH, Qi R, Liu S, Kroh A, Yao G, Perry K, Rummel A, Jin R. 2017. The hypothetical protein P47 of *Clostridium botulinum* E1 strain Beluga has a structural topology similar to bactericidal/permeability-increasing protein. Toxicon. pii: S0041-0101(17)30309-4.

16. Lawson, Paul. 2016a. The taxonomy of the genus *Clostridium*: Current status and future perspectives. Conference: The 7th National Conference of Microbial Resources & the International Symposium on Microbial Systematics and Taxonomy, At Hangzhou, China, Microbioogy China 43(5): 1070–1074

17. Lawson PA, Citron DM, Tyrrell KL, Finegold SM. 2016b. Reclassification of *Clostridium difficile* as *Clostridioides difficile* (Hall and O’Toole 1935) Prévot 1938. Anaerobe. 40:95–9.

18. Licona-Cassani C, Steen JA, Zaragoza NE, Moonen G, Moutafis G, Hodson MP, Power J, Nielsen LK, Marcellin E. 2016. Tetanus toxin production is triggered by the transition from amino acid consumption to peptides. Anaerobe. 41:113–124.

19. Liou JS, Balkwill DL, Drake GR, Tanner RS. 2005. *Clostridium carboxidivorans* sp. nov., a solvent-producing *Clostridium* isolated from an agricultural settling lagoon, and reclassification of the acetogen *Clostridium scatologenes* strain SL1 as *Clostridium drakei* sp. nov. Int J Syst Evol Microbiol. 55(Pt 5):2085–91.

20. Nguyen LT, Schmidt HA, von Haeseler A, Minh BQ. 2015. IQ-TREE: A fast and effective stochastic algorithm for estimating maximum likelihood phylogenies. Mol. Biol. Evol., 32:268–274.

21. Parker CT, Tindall BJ, and Garrity GM. 2015. International Code of Nomenclature of Prokaryotes. Int J Syst Evol Microbiol. doi: 10.1099/ijsem.0.000778.

22. Petit L, Gibert M, and Popoff MR. (1999). *Clostridium perfringens*: toxinotype and genotype. Trends Microbiol. 7(3): 104–110.

23. Rainey F, Tanner R, and Wiegel J. 2006 Family Clostridiaceae, in “the Prokaryotes: A Handbook on the Biology of Bacteria: Vol. 4: Bacteria: Firmicutes, Cyanobacteria”, 3rd edn. release 3.20., New York: Springer-Verlag, pp. 4:654–678 (DOI: 10.1007/0-387-30744-3_20.

24. Schniete JK, Cruz-Morales P, Selem-Mojica N, Fernández-Martínez LT, Hunter IS, Barona-Gómez F, Hoskisson PA. 2018. Expanding primary metabolism helps generate the metabolic robustness to facilitate antibiotic biosynthesis in *Streptomyces*. mBio 9:e02283–17.

25. Shiratori H, Sasaya K, Ohiwa H, Ikeno H, Ayame S, Kataoka N, Miya A, Beppu T, Ueda K. 2009. *Clostridium clariflavum* sp. nov. and *Clostridium caenicola* sp. nov., moderately thermophilic, cellulose-/cellobiose-digesting bacteria isolated from methanogenic sludge. Int J Syst Evol Microbiol. 59(Pt 7):1764–70.

1. Skarin H, and Segerman B. 2011. Horizontal gene transfer of toxin genes in *Clostridium botulinum*: Involvement of mobile elements and plasmids. Mob Genet Elements. 1(3): 213– 215.

2. Slobodkina GB, Kolganova TV, Tourova TP, Kostrikina NA, Jeanthon C, Bonch-Osmolovskaya EA, Slobodkin AI.2008. *Clostridium tepidiprofundi* sp. nov., a moderately thermophilic bacterium from a deep-sea hydrothermal vent. Int J Syst Evol Microbiol. 58(Pt 4):852–5.

3. Tamburini E, Daly S, Steiner U, Vandini C, Mastromei G. 2001. *Clostridium felsineum* and *Clostridium acetobutylicum* are two distinct species that are phylogenetically closely related. Int J Syst Evol Microbiol. 51(Pt 3):963–6.

4. Udaondo Z, Duque E, Ramos JL. 2017. The pangenome of the genus *Clostridium*. Environ Microbiol. 19(7):2588–2603.

5. Vos RA, Balhoff JP, Caravas JA, Holder MT, Lapp H. Maddison WP, Midford P E, Priyam A, Sukumaran J, Xia X and Stoltzfus A. 2012. NeXML: rich, extensible, and verifiable representation of comparative data and metadata. Systematic Biology 61(4): 675–689

6. Warren YA, Tyrrell KL, Citron DM, Goldstein EJ. 2006. *Clostridium aldenense* sp. nov. and *Clostridium citroniae* sp. nov. isolated from human clinical infections. J Clin Microbiol. 44(7):2416–22.

7. Weigand MR, Pena-Gonzalez A, Shirey TB, Broeker RG, Ishaq MK, Konstantinidis KT, Raphael BH. 2015. Implications of Genome-Based Discrimination between *Clostridium botulinum* Group I and *Clostridium sporogenes* Strains for Bacterial Taxonomy. Appl Environ Microbiol. 81(16):5420–9

8. Yutin N, Galperin MY. 2013. A genomic update on clostridial phylogeny: Gram-negative spore formers and other misplaced clostridia. Environ Microbiol. 15(10):2631–41.

