## Supplementary material for "Revisiting the evolution and taxonomy of Clostridia, a phylogenomic update"

Pablo Cruz-Morales^1,a^, Camila A. Orellana^1^, George Moutafis^2^, Glenn Moonen^2^, Gonzalo Rincon^2^, Lars K Nielsen^1^ and Esteban Marcellin^1^

^1^Australian Institute for Bioengineering and Nanotechnology, The University of Queensland

^2^ Zoetis, 45 Poplar Rd, Parkville, Victoria Australia 3052

^a^ Present address: Joint BioEnergy Institute, Lawrence Berkeley National Laboratory and Centro de Biotecnología FEMSA, Instituo Tecnologico y de Estudios Superiores de Monterrey

**Table S1. Genomes included in the analysis**

| **Organism name** | **Clostridial cluster** | **Contigs** | **Length** | **N50** | **L50** |
| --- | --- | --- | --- | --- | --- |
| Clostridium_acetobutylicum_ATCC_824_2 | I | 2 | 4132880 | 3940880 | 1 |
| Clostridium_acetobutylicum_DSM_1731_61 | I | 3 | 4145581 | 3942462 | 1 |
| Clostridium_acetobutylicum_DSM_1732_527 | I | 55 | 4091215 | 270881 | 6 |
| Clostridium_acetobutylicum_EA_2018_57 | I | 2 | 4132226 | 3940230 | 1 |
| Clostridium_acetobutylicum_GXAS18_1_231 | I | 49 | 3796049 | 325351 | 4 |
| Clostridium_acetobutylicum_NCCB_24020_525 | I | 20 | 4098731 | 759218 | 2 |
| Clostridium_akagii_DSM_12554_164 | I | 50 | 4589511 | 266036 | 6 |
| Clostridium_algidicarnis_B3_167 | I | 1 | 3060291 | 3060291 | 1 |
| Clostridium_arbusti_SL206_70 | I | 243 | 3970278 | 31124 | 40 |
| Clostridium_argentinense_CDC_2741_240 | I | 20 | 4742562 | 418119 | 4 |
| Clostridium_aurantibutyricum_DSM_793_518 | I | 221 | 4922827 | 51056 | 31 |
| Clostridium_baratii_2789STDY5834907_337 | I | 50 | 3141705 | 1732372 | 1 |
| Clostridium_baratii_2789STDY5834956_336 | I | 30 | 3086202 | 440129 | 2 |
| Clostridium_baratii_771_14_262 | I | 39 | 3173174 | 211064 | 5 |
| Clostridium_baratii_CDC51267_505 | I | 2 | 3211717 | 3091050 | 1 |
| Clostridium_baratii_str._Sullivan_233 | I | 2 | 3338630 | 3153266 | 1 |
| Clostridium_baratii_XCM_325 | I | 28 | 3087740 | 857756 | 2 |
| Clostridium_beijerinckii_4J9_513 | I | 162 | 5888124 | 89920 | 20 |
| Clostridium_beijerinckii_ATCC_35702_SA_1_224 | I | 1 | 5999050 | 5999050 | 1 |
| Clostridium_beijerinckii_ATCC_39058_512 | I | 302 | 5953339 | 49706 | 37 |
| Clostridium_beijerinckii_BAS_B2_530 | I | 245 | 5982920 | 58724 | 32 |
| Clostridium_beijerinckii_BAS_B3_I_124_509 | I | 1 | 6123550 | 6123550 | 1 |
| Clostridium_beijerinckii_BGS1_439 | I | 105 | 5880896 | 195648 | 10 |
| Clostridium_beijerinckii_DSM_53_516 | I | 346 | 5773247 | 40593 | 42 |
| Clostridium_beijerinckii_DSM_791_529 | I | 264 | 5781472 | 43059 | 41 |
| Clostridium_beijerinckii_G117_75 | I | 89 | 5811816 | 172488 | 14 |
| Clostridium_beijerinckii_HUN142_159 | I | 53 | 6106710 | 236200 | 8 |
| Clostridium_beijerinckii_NCIMB_14988_251 | I | 1 | 6485394 | 6485394 | 1 |
| Clostridium_beijerinckii_NCIMB_8052_8 | I | 1 | 6000632 | 6000632 | 1 |
| Clostridium_beijerinckii_NCP_260_520 | I | 242 | 5968330 | 59545 | 30 |
| Clostridium_beijerinckii_NRRL_B_528_526 | I | 233 | 6255488 | 70635 | 27 |
| Clostridium_beijerinckii_NRRL_B_591_521 | I | 358 | 5874824 | 39987 | 43 |
| Clostridium_beijerinckii_NRRL_B_593_522 | I | 305 | 6156662 | 46401 | 38 |
| Clostridium_beijerinckii_NRRL_B_596_528 | I | 393 | 6220133 | 38644 | 47 |
| Clostridium_beijerinckii_NRRL_B_598_138 | I | 1 | 6186879 | 6186879 | 1 |
| Clostridium_botulinum_111_249 | I | 1 | 3901300 | 3901300 | 1 |
| Clostridium_botulinum_12LNR10_CD_302 | I | 132 | 3041748 | 39896 | 22 |
| Clostridium_botulinum_12LNR13_CD_304 | I | 140 | 3075461 | 41782 | 24 |
| Clostridium_botulinum_12LNRI_CD_299 | I | 131 | 3005803 | 41798 | 21 |
| Clostridium_botulinum_202F_232 | I | 2 | 3914604 | 3874462 | 1 |
| Clostridium_botulinum_211_366 | I | 42 | 3712985 | 404531 | 4 |
| Clostridium_botulinum_246 | I | 1 | 3611897 | 3611897 | 1 |
| Clostridium_botulinum_277_00_250 | I | 90 | 3938115 | 133135 | 10 |
| Clostridium_botulinum_29401_CD_303 | I | 112 | 3046146 | 57518 | 18 |
| Clostridium_botulinum_38028_CD_301 | I | 104 | 3118863 | 48579 | 19 |
| Clostridium_botulinum_43243_CD_298 | I | 111 | 3005167 | 57521 | 16 |
| Clostridium_botulinum_47295_DC_309 | I | 108 | 3178512 | 51664 | 20 |
| Clostridium_botulinum_48212_CD_300 | I | 75 | 3005337 | 68539 | 14 |
| Clostridium_botulinum_49511_CD_307 | I | 85 | 3088897 | 64109 | 12 |
| Clostridium_botulinum_50867_CD_306 | I | 134 | 3077301 | 36158 | 22 |
| Clostridium_botulinum_51714_DC_308 | I | 101 | 3174715 | 59200 | 19 |
| Clostridium_botulinum_55741_CD_305 | I | 103 | 3040725 | 57521 | 17 |
| Clostridium_botulinum_58272_CD_313 | I | 128 | 3074857 | 40020 | 22 |
| Clostridium_botulinum_58752_CD_314 | I | 114 | 2825871 | 41221 | 20 |
| Clostridium_botulinum_69285_CD_310 | I | 146 | 2937195 | 61722 | 15 |
| Clostridium_botulinum_713_CBOT_288 | I | 191 | 3551054 | 41974 | 25 |
| Clostridium_botulinum_71840_CD_311 | I | 145 | 3001250 | 40406 | 22 |
| Clostridium_botulinum_A_str._ATCC_19397_9 | I | 1 | 3863450 | 3863450 | 1 |
| Clostridium_botulinum_A_str._ATCC_3502_19 | I | 2 | 3903260 | 3886916 | 1 |
| Clostridium_botulinum_A_str._Hall_10 | I | 1 | 3760560 | 3760560 | 1 |
| Clostridium_botulinum_A2_117_173 | I | 11 | 3808262 | 2752809 | 1 |
| Clostridium_botulinum_A2_str._Kyoto_18 | I | 1 | 4155278 | 4155278 | 1 |
| Clostridium_botulinum_A2B3_87_171 | I | 13 | 4168550 | 2767020 | 1 |
| Clostridium_botulinum_A2B7_92_172 | I | 14 | 4057812 | 698075 | 2 |
| Clostridium_botulinum_A3_str._Loch_Maree_13 | I | 2 | 4259691 | 3992906 | 1 |
| Clostridium_botulinum_AM1195_389 | I | 31 | 4271150 | 299090 | 6 |
| Clostridium_botulinum_AM1295_390 | I | 34 | 3900045 | 410269 | 3 |
| Clostridium_botulinum_AM370_391 | I | 31 | 4269660 | 398258 | 4 |
| Clostridium_botulinum_AM553_392 | I | 30 | 4308056 | 685346 | 3 |
| Clostridium_botulinum_An436_367 | I | 70 | 4201460 | 117448 | 11 |
| Clostridium_botulinum_ATCC_17786_322 | I | 61 | 3951940 | 193297 | 7 |
| Clostridium_botulinum_ATCC_17843_316 | I | 24 | 3906754 | 2083105 | 1 |
| Clostridium_botulinum_ATCC_17843_B5_365 | I | 79 | 3898546 | 116115 | 10 |
| Clostridium_botulinum_ATCC_17862_319 | I | 56 | 3907623 | 227875 | 6 |
| Clostridium_botulinum_ATCC_23387_317 | I | 29 | 3817674 | 790194 | 2 |
| Clostridium_botulinum_ATCC_25763_323 | I | 28 | 3889092 | 720423 | 3 |
| Clostridium_botulinum_ATCC_449_369 | I | 65 | 3777532 | 137365 | 10 |
| Clostridium_botulinum_ATCC_7949_320 | I | 35 | 3909427 | 612792 | 3 |
| Clostridium_botulinum_ATCC_9564_318 | I | 57 | 3813606 | 201673 | 7 |
| Clostridium_botulinum_B_str_Eklund_17B_NRP_14 | I | 2 | 3847969 | 3800327 | 1 |
| Clostridium_botulinum_B_str._Osaka05_137 | I | 3 | 4408349 | 4004744 | 1 |
| Clostridium_botulinum_B1_str._Okra_12 | I | 2 | 4107013 | 3958233 | 1 |
| Clostridium_botulinum_B2_128_174 | I | 10 | 3844467 | 2786970 | 1 |
| Clostridium_botulinum_B2_267_175 | I | 12 | 3903580 | 2837736 | 1 |
| Clostridium_botulinum_B2_275_170 | I | 13 | 3978188 | 2203691 | 1 |
| Clostridium_botulinum_B2_331_169 | I | 10 | 3809103 | 2732590 | 1 |
| Clostridium_botulinum_B2_433_177 | I | 12 | 4124526 | 2803320 | 1 |
| Clostridium_botulinum_B2_450_252 | I | 10 | 4320669 | 2916972 | 1 |
| Clostridium_botulinum_Ba4_657_16 | I | 3 | 4257769 | 3977794 | 1 |
| Clostridium_botulinum_Bf_39 | I | 70 | 4217754 | 170315 | 9 |
| Clostridium_botulinum_BKT015925_60 | I | 6 | 3207592 | 2773157 | 1 |
| Clostridium_botulinum_BKT028387_59 | I | 237 | 2833823 | 29150 | 33 |
| Clostridium_botulinum_C_D_str._BKT12695_168 | I | 89 | 2754780 | 63795 | 14 |
| Clostridium_botulinum_C_D_str._BKT2873_184 | I | 159 | 3165510 | 77101 | 14 |
| Clostridium_botulinum_C_D_str._BKT75002_185 | I | 121 | 3138765 | 77530 | 14 |
| Clostridium_botulinum_C_D_str._It1_188 | I | 77 | 2499706 | 59587 | 15 |
| Clostridium_botulinum_C_D_str._Sp77_187 | I | 148 | 3057314 | 40000 | 22 |
| Clostridium_botulinum_C_str._Eklund_40 | I | 76 | 2961186 | 145667 | 6 |
| Clostridium_botulinum_CB11_1_1_90 | I | 171 | 3823307 | 43401 | 26 |
| Clostridium_botulinum_CDC_1436_242 | I | 2 | 4365669 | 4089683 | 1 |
| Clostridium_botulinum_CDC_1632_470 | I | 1 | 4393047 | 4393047 | 1 |
| Clostridium_botulinum_CDC_1744_364 | I | 71 | 3959495 | 142985 | 8 |
| Clostridium_botulinum_CDC_5247_384 | I | 51 | 3839610 | 203210 | 5 |
| Clostridium_botulinum_CDC_53174_471 | I | 1 | 3867627 | 3867627 | 1 |
| Clostridium_botulinum_CDC_67190_468 | I | 3 | 4020063 | 3954777 | 1 |
| Clostridium_botulinum_CDC_69094_467 | I | 1 | 4089027 | 4089027 | 1 |
| Clostridium_botulinum_CDC_69096_503 | I | 26 | 4252578 | 3982791 | 1 |
| Clostridium_botulinum_CDC_795_381 | I | 97 | 3920281 | 96784 | 13 |
| Clostridium_botulinum_CDC28023_195 | I | 341 | 3966737 | 22262 | 51 |
| Clostridium_botulinum_CDC48719_194 | I | 310 | 3977994 | 26132 | 44 |
| Clostridium_botulinum_CDC66177_81 | I | 119 | 3852437 | 86296 | 15 |
| Clostridium_botulinum_CFSAN024410_230 | I | 131 | 4005128 | 59804 | 22 |
| Clostridium_botulinum_D_CCUG_7971_229 | I | 111 | 2808469 | 52759 | 17 |
| Clostridium_botulinum_D_str._16868_186 | I | 129 | 3083628 | 95198 | 12 |
| Clostridium_botulinum_DB_2_211 | I | 150 | 3915341 | 288306 | 5 |
| Clostridium_botulinum_E1_str._BoNT_E_Beluga_48 | I | 6 | 3999201 | 3863095 | 1 |
| Clostridium_botulinum_E3_str._Alaska_E43_15 | I | 1 | 3659644 | 3659644 | 1 |
| Clostridium_botulinum_F_357_176 | I | 15 | 3832122 | 957457 | 2 |
| Clostridium_botulinum_F_str._230613_20 | I | 2 | 4010614 | 3993083 | 1 |
| Clostridium_botulinum_F_str._Langeland_11 | I | 2 | 4012918 | 3995387 | 1 |
| Clostridium_botulinum_H04402_065_71 | I | 1 | 3919740 | 3919740 | 1 |
| Clostridium_botulinum_Hall_183_370 | I | 99 | 4002127 | 78163 | 16 |
| Clostridium_botulinum_Hall_4834_371 | I | 86 | 4025192 | 97924 | 11 |
| Clostridium_botulinum_Hall_80_378 | I | 65 | 3812410 | 175053 | 9 |
| Clostridium_botulinum_Hall_8388A_374 | I | 55 | 3909258 | 191646 | 7 |
| Clostridium_botulinum_Hall_8857Ab_372 | I | 84 | 3971034 | 101664 | 11 |
| Clostridium_botulinum_K15_393 | I | 114 | 3997935 | 100195 | 13 |
| Clostridium_botulinum_K3_394 | I | 218 | 3850229 | 221625 | 4 |
| Clostridium_botulinum_KAPB_3_210 | I | 128 | 3871084 | 726694 | 3 |
| Clostridium_botulinum_KF_Meyer_126_373 | I | 55 | 3891395 | 199059 | 7 |
| Clostridium_botulinum_KF_Meyer_33_380 | I | 98 | 3896775 | 72415 | 15 |
| Clostridium_botulinum_L_572_383 | I | 82 | 3766552 | 99892 | 11 |
| Clostridium_botulinum_LNC5_DC_312 | I | 83 | 2894028 | 55876 | 15 |
| Clostridium_botulinum_Mauritius_395 | I | 84 | 3869437 | 100079 | 13 |
| Clostridium_botulinum_McClung_844_379 | I | 48 | 3856611 | 219996 | 6 |
| Clostridium_botulinum_NCTC_2916_38 | I | 49 | 4031357 | 433501 | 3 |
| Clostridium_botulinum_NCTC_8550_247 | I | 1 | 3611898 | 3611898 | 1 |
| Clostridium_botulinum_Prevot_594_241 | I | 2 | 4334551 | 4077214 | 1 |
| Clostridium_botulinum_Prevot_697B_377 | I | 34 | 3791338 | 344362 | 4 |
| Clostridium_botulinum_Prevot_910_375 | I | 74 | 3942683 | 116912 | 10 |
| Clostridium_botulinum_Prevot_Dewping_376 | I | 42 | 3880168 | 252959 | 5 |
| Clostridium_botulinum_Prevot_Ped_1_385 | I | 81 | 3791027 | 125071 | 10 |
| Clostridium_botulinum_Prevot_R81_3A_386 | I | 64 | 3783752 | 200588 | 6 |
| Clostridium_botulinum_SU0634_420 | I | 43 | 3882852 | 201043 | 6 |
| Clostridium_botulinum_SU0635W_422 | I | 137 | 3945914 | 64297 | 16 |
| Clostridium_botulinum_SU0729_421 | I | 72 | 3929760 | 270392 | 6 |
| Clostridium_botulinum_SU0801_416 | I | 124 | 4210622 | 117792 | 10 |
| Clostridium_botulinum_SU0807 | I | 68 | 4091988 | 249852 | 6 |
| Clostridium_botulinum_SU0945_415 | I | 272 | 4534970 | 140479 | 11 |
| Clostridium_botulinum_SU0994_419 | I | 51 | 4296977 | 372385 | 5 |
| Clostridium_botulinum_SU0998_418 | I | 61 | 4136384 | 274519 | 5 |
| Clostridium_botulinum_SU1054_409 | I | 99 | 4311895 | 274519 | 5 |
| Clostridium_botulinum_SU1064_411 | I | 139 | 4368910 | 424584 | 4 |
| Clostridium_botulinum_SU1072_410 | I | 66 | 4281988 | 274519 | 4 |
| Clostridium_botulinum_SU1074_413 | I | 61 | 4055427 | 429106 | 4 |
| Clostridium_botulinum_SU1112_414 | I | 46 | 3929155 | 460210 | 3 |
| Clostridium_botulinum_SU1169_412 | I | 204 | 4495542 | 197058 | 8 |
| Clostridium_botulinum_SU1259_407 | I | 174 | 4175404 | 723257 | 2 |
| Clostridium_botulinum_SU1274_401 | I | 50 | 4230750 | 274519 | 6 |
| Clostridium_botulinum_SU1275_404 | I | 280 | 4305633 | 150850 | 7 |
| Clostridium_botulinum_SU1304_406 | I | 141 | 4306760 | 222901 | 6 |
| Clostridium_botulinum_SU1306_399 | I | 66 | 4267307 | 146794 | 11 |
| Clostridium_botulinum_SU1575NT_400 | I | 38 | 3873577 | 357116 | 4 |
| Clostridium_botulinum_SU1887_403 | I | 67 | 4010957 | 238528 | 4 |
| Clostridium_botulinum_SU1891_408 | I | 223 | 4476519 | 277699 | 5 |
| Clostridium_botulinum_SU1917_402 | I | 61 | 4274979 | 215880 | 5 |
| Clostridium_botulinum_SU1934_405 | I | 38 | 3990632 | 529942 | 2 |
| Clostridium_botulinum_SU1937_398 | I | 85 | 4285737 | 214161 | 5 |
| Clostridium_botulinum_type_C_BOTC | I | 135 | 2617336 | 145081 | 6 |
| Clostridium_botulinum_type_D_BOTD | I | 259 | 3059863 | 55039 | 18 |
| Clostridium_botulinum_V891_58 | I | 114 | 3172457 | 68878 | 16 |
| Clostridium_botulinum_VPI_7124_368 | I | 80 | 3864510 | 77273 | 16 |
| Clostridium_botulinum_Walls_8G_280 | I | 27 | 3978537 | 835326 | 2 |
| Clostridium_butyricum_5521_41 | I | 123 | 4540699 | 81014 | 17 |
| Clostridium_butyricum_60E.3_105 | I | 10 | 4644398 | 2372918 | 1 |
| Clostridium_butyricum_AGR2140_121 | I | 39 | 4550822 | 237318 | 7 |
| Clostridium_butyricum_CDC_51208_469 | I | 3 | 4639914 | 3809831 | 1 |
| Clostridium_butyricum_CWBI1009_255 | I | 340 | 4491619 | 26889 | 51 |
| Clostridium_butyricum_DKU_01_91 | I | 79 | 4519722 | 108221 | 13 |
| Clostridium_butyricum_DSM_10702_116 | I | 207 | 4596811 | 83186 | 17 |
| Clostridium_butyricum_E4_str._BoNT_E_BL5262_52 | I | 13 | 4758422 | 757653 | 2 |
| Clostridium_butyricum_HM_68_253 | I | 2 | 4604758 | 3835983 | 1 |
| Clostridium_butyricum_JKY6D1_355 | I | 3 | 4618327 | 3819894 | 1 |
| Clostridium_butyricum_KNU_L09_352 | I | 2 | 4627894 | 3824894 | 1 |
| Clostridium_butyricum_TOA_434 | I | 3 | 4597202 | 3794139 | 1 |
| Clostridium_cadaveris_AGR2141_120 | I | 42 | 3542160 | 141803 | 8 |
| Clostridium_cadaveris_NLAE_zl_G419_543 | I | 52 | 3532192 | 120460 | 10 |
| Clostridium_carboxidivorans_P7_284 | I | 2 | 5752782 | 5732880 | 1 |
| Clostridium_cavendishii_DSM_21758_555 | I | 56 | 4987666 | 284556 | 7 |
| Clostridium_chauvoei_JF4335_1112 | I | 1 | 2887451 | 2887451 | 1 |
| Clostridium_chauvoeiCH3_CHAUPV | I | 309 | 2892977 | 72373 | 13 |
| Clostridium_chauvoeiCN3796_CHAU3796 | I | 292 | 2935638 | 79097 | 11 |
| Clostridium_colicanis_209318_97 | I | 10 | 3505620 | 2922022 | 1 |
| Clostridium_collagenovorans_DSM_3089_553 | I | 30 | 3482404 | 417404 | 3 |
| Clostridium_coskatii_PS02_430 | I | 85 | 4565108 | 130355 | 12 |
| Clostridium_coskatii_PTA_10522_437 | I | 112 | 4538837 | 90816 | 16 |
| Clostridium_disporicum_2789STDY5608827_335 | I | 110 | 3739749 | 84850 | 15 |
| Clostridium_disporicum_2789STDY5834855_330 | I | 119 | 3804830 | 80327 | 18 |
| Clostridium_disporicum_2789STDY5834856_333 | I | 57 | 3449564 | 144642 | 9 |
| Clostridium_drakei_SL1_160 | I | 122 | 5578774 | 151002 | 12 |
| Clostridium_estertheticum_subsp._estertheticum_DSM_8809_466 | I | 2 | 4785613 | 4760574 | 1 |
| Clostridium_fallax_DSM_2631_549 | I | 64 | 2747337 | 94344 | 9 |
| Clostridium_felsineum_DSM_794_524 | I | 100 | 5178654 | 177705 | 10 |
| Clostridium_haemolyticum_3629_HAEMOLYT | I | 285 | 2650828 | 55585 | 16 |
| Clostridium_haemolyticum_NCTC_8350_225 | I | 234 | 2465609 | 42678 | 18 |
| Clostridium_haemolyticum_NCTC_9693_180 | I | 125 | 2607656 | 55585 | 15 |
| Clostridium_hydrogeniformans_DSM_21757_163 | I | 31 | 4101506 | 314904 | 4 |
| Clostridium_intestinale_DSM_6191_554 | I | 33 | 4600598 | 368167 | 5 |
| Clostridium_intestinale_URNW_131 | I | 3 | 4677668 | 4667789 | 1 |
| Clostridium_kluyveri_DSM_555_7 | I | 2 | 4023800 | 3964618 | 1 |
| Clostridium_kluyveri_JZZ_472 | I | 2 | 4512934 | 4454353 | 1 |
| Clostridium_kluyveri_NBRC_12016_4 | I | 2 | 3955303 | 3896121 | 1 |
| Clostridium_ljungdahlii_DSM_13528_21 | I | 1 | 4630065 | 4630065 | 1 |
| Clostridium_ljungdahlii_DSM_13528_PETC_432 | I | 56 | 4572618 | 200112 | 8 |
| Clostridium_ljungdahlii_ERI_2_431 | I | 65 | 4363917 | 161324 | 7 |
| Clostridium_magnum_DSM_2767_429 | I | 25 | 6634930 | 740872 | 3 |
| Clostridium_novyi_A_str._4540_191 | I | 81 | 2500001 | 62812 | 12 |
| Clostridium_novyi_A_str._4552_228 | I | 119 | 2798065 | 47012 | 21 |
| Clostridium_novyi_A_str._4570_226 | I | 69 | 2324088 | 71398 | 9 |
| Clostridium_novyi_A_str._BKT29909_190 | I | 95 | 2463931 | 45649 | 17 |
| Clostridium_novyi_A_str._GD211209_189 | I | 104 | 2459310 | 38497 | 19 |
| Clostridium_novyi_A_str._NCTC_538_192 | I | 69 | 2519932 | 63662 | 14 |
| Clostridium_novyi_B_str._ATCC_27606_181 | I | 168 | 2613094 | 46394 | 20 |
| Clostridium_novyi_NT_6 | I | 1 | 2547720 | 2547720 | 1 |
| Clostridium_novyi_Type_BT46_oxer_NOVYIPV | I | 245 | 2606506 | 62864 | 15 |
| Clostridium_paraputrificum_2789STDY5834857_331 | I | 25 | 3597627 | 348122 | 4 |
| Clostridium_paraputrificum_373_A1_438 | I | 41 | 3488595 | 259466 | 5 |
| Clostridium_paraputrificum_AGR2156_119 | I | 30 | 3561289 | 320228 | 4 |
| Clostridium_pasteurianum_BC1_112 | I | 2 | 5044100 | 4990707 | 1 |
| Clostridium_pasteurianum_DSM_525_ATCC_6013_464 | I | 1 | 4352852 | 4352852 | 1 |
| Clostridium_pasteurianum_GL11_443 | I | 2 | 4677181 | 4485768 | 1 |
| Clostridium_pasteurianum_M150B_465 | I | 1 | 4351863 | 4351863 | 1 |
| Clostridium_perfringens_1207_CPER_287 | I | 95 | 3207161 | 85634 | 11 |
| Clostridium_perfringens_2789STDY5608889_338 | I | 48 | 3321076 | 2127284 | 1 |
| Clostridium_perfringens_ATCC_13124_5 | I | 1 | 3256683 | 3256683 | 1 |
| Clostridium_perfringens_B_str._ATCC_3626_43 | I | 98 | 3896305 | 88742 | 12 |
| Clostridium_perfringens_C_str._JGS1495_42 | I | 84 | 3661329 | 117588 | 10 |
| Clostridium_perfringens_CP4_343 | I | 98 | 3642209 | 83876 | 13 |
| Clostridium_perfringens_CPE_str._F4969_45 | I | 74 | 3510272 | 96499 | 10 |
| Clostridium_perfringens_E_str._JGS1987_44 | I | 101 | 4127102 | 88895 | 14 |
| Clostridium_perfringens_F262_72 | I | 14 | 3468406 | 3333039 | 1 |
| Clostridium_perfringens_FORC_003_aquarium_water_326 | I | 2 | 3395109 | 3338532 | 1 |
| Clostridium_perfringens_FORC_025_462 | I | 1 | 3343822 | 3343822 | 1 |
| Clostridium_perfringens_JFP718_473 | I | 56 | 3652220 | 197638 | 7 |
| Clostridium_perfringens_JFP727_476 | I | 47 | 3624033 | 151112 | 8 |
| Clostridium_perfringens_JFP728_475 | I | 85 | 3579626 | 83659 | 15 |
| Clostridium_perfringens_JFP771_481 | I | 81 | 3495308 | 118182 | 10 |
| Clostridium_perfringens_JFP774_474 | I | 56 | 3548595 | 142704 | 7 |
| Clostridium_perfringens_JFP795_477 | I | 67 | 3578912 | 114607 | 12 |
| Clostridium_perfringens_JFP796_480 | I | 114 | 3601148 | 75633 | 17 |
| Clostridium_perfringens_JFP801_478 | I | 59 | 3580610 | 143793 | 8 |
| Clostridium_perfringens_JFP804_479 | I | 117 | 3625968 | 71788 | 15 |
| Clostridium_perfringens_JFP810_482 | I | 95 | 3830406 | 161738 | 8 |
| Clostridium_perfringens_JFP826_483 | I | 70 | 3660408 | 105999 | 10 |
| Clostridium_perfringens_JFP828_484 | I | 54 | 3564370 | 125860 | 10 |
| Clostridium_perfringens_JFP829_485 | I | 108 | 3639686 | 71458 | 18 |
| Clostridium_perfringens_JFP833_486 | I | 96 | 3521770 | 84324 | 13 |
| Clostridium_perfringens_JFP834_487 | I | 121 | 3549140 | 96309 | 13 |
| Clostridium_perfringens_JFP836_488 | I | 51 | 3599532 | 163794 | 7 |
| Clostridium_perfringens_JFP914_489 | I | 79 | 3819232 | 104224 | 13 |
| Clostridium_perfringens_JFP916_490 | I | 67 | 3660641 | 160443 | 9 |
| Clostridium_perfringens_JFP921_491 | I | 78 | 3601767 | 100577 | 12 |
| Clostridium_perfringens_JFP922_492 | I | 76 | 3612099 | 186517 | 7 |
| Clostridium_perfringens_JFP923_493 | I | 48 | 3589715 | 212576 | 6 |
| Clostridium_perfringens_JFP941_494 | I | 64 | 3594640 | 109035 | 11 |
| Clostridium_perfringens_JFP961_495 | I | 66 | 3617219 | 138753 | 9 |
| Clostridium_perfringens_JFP978_496 | I | 68 | 3585277 | 106630 | 12 |
| Clostridium_perfringens_JFP980_497 | I | 67 | 3654588 | 108867 | 11 |
| Clostridium_perfringens_JFP981_498 | I | 68 | 3580367 | 131646 | 10 |
| Clostridium_perfringens_JFP982_499 | I | 62 | 3670557 | 137154 | 8 |
| Clostridium_perfringens_JFP983_500 | I | 46 | 3657522 | 204880 | 7 |
| Clostridium_perfringens_JFP986_502 | I | 58 | 3669049 | 158758 | 9 |
| Clostridium_perfringens_JFP99_501 | I | 70 | 3657868 | 174400 | 8 |
| Clostridium_perfringens_JJC_140 | I | 69 | 3259329 | 98246 | 9 |
| Clostridium_perfringens_JP55_423 | I | 6 | 3571229 | 3347300 | 1 |
| Clostridium_perfringens_JP838_424 | I | 5 | 4070811 | 3530414 | 1 |
| Clostridium_perfringens_MJR7757A_361 | I | 228 | 3590266 | 51972 | 22 |
| Clostridium_perfringens_NCTC_8239_str._NCTC_8239_46 | I | 55 | 3324319 | 134604 | 6 |
| Clostridium_perfringens_str._13_3 | I | 2 | 3085740 | 3031430 | 1 |
| Clostridium_perfringens_Type_A_PERACSIR | I | 228 | 3483931 | 226748 | 4 |
| Clostridium_perfringens_Type_A1491_PERA1491 | I | 140 | 3279841 | 2055711 | 1 |
| Clostridium_perfringens_Type_B1240_PERB1240 | I | 122 | 3547357 | 261573 | 3 |
| Clostridium_perfringens_Type_B3424_PERB3424 | I | 243 | 3732983 | 410377 | 3 |
| Clostridium_perfringens_Type_C883_PERC883 | I | 282 | 3683980 | 291793 | 4 |
| Clostridium_perfringens_Type_D3688_PERD3683 | I | 166 | 3687986 | 238842 | 4 |
| Clostridium_perfringens_Type_DSB170D_PERDPV | I | 115 | 3495957 | 237771 | 4 |
| Clostridium_perfringens_WAL_14572_68 | I | 20 | 3466039 | 2084089 | 1 |
| Clostridium_proteolyticum_DSM_3090_557 | I | 45 | 2818024 | 121571 | 8 |
| Clostridium_puniceum_DSM_2619_523 | I | 245 | 6082167 | 68127 | 31 |
| Clostridium_roseum_DSM_6424_515 | I | 262 | 4944863 | 49535 | 32 |
| Clostridium_roseum_DSM_7320_517 | I | 124 | 5067725 | 86225 | 18 |
| Clostridium_saccharobutylicum_BAS_B3_SW_136_508 | I | 1 | 5108304 | 5108304 | 1 |
| Clostridium_saccharobutylicum_DSM_13864_133 | I | 1 | 5107814 | 5107814 | 1 |
| Clostridium_saccharobutylicum_L1_8_514 | I | 16 | 5173344 | 482513 | 4 |
| Clostridium_saccharobutylicum_NCP_165_519 | I | 142 | 4900327 | 83530 | 16 |
| Clostridium_saccharobutylicum_NCP_195_511 | I | 1 | 5108176 | 5108176 | 1 |
| Clostridium_saccharobutylicum_NCP_200_506 | I | 1 | 5108287 | 5108287 | 1 |
| Clostridium_saccharobutylicum_NCP_258_510 | I | 1 | 4950933 | 4950933 | 1 |
| Clostridium_saccharoperbutylacetonicum_N1_4HMT_87 | I | 2 | 6666445 | 6530257 | 1 |
| Clostridium_saccharoperbutylacetonicum_N1_4HMT_ATCC_27021_86 | I | 210 | 6569628 | 78354 | 30 |
| Clostridium_saccharoperbutylacetonicum_N1_504_507 | I | 2 | 6219394 | 6216458 | 1 |
| Clostridium_sartagoforme_AAU1_114 | I | 323 | 3983371 | 25206 | 45 |
| Clostridium_scatologenes_ATCC_25775_278 | I | 1 | 5749410 | 5749410 | 1 |
| Clostridium_septicum_P1044_537 | I | 79 | 3298970 | 71267 | 16 |
| Clostridium_septicum38_858V_SEPTPV | I | 342 | 3373031 | 50613 | 21 |
| Clostridium_septicumCN3204_SEP3204B | I | 218 | 3344920 | 52620 | 20 |
| Clostridium_septicumCN368_SEPT368 | I | 243 | 3308673 | 44576 | 21 |
| Clostridium_sporogenes_1961_2_345 | I | 40 | 4092065 | 213496 | 5 |
| Clostridium_sporogenes_1990_344 | I | 59 | 4163429 | 187341 | 7 |
| Clostridium_sporogenes_2007_351 | I | 33 | 4170948 | 321923 | 4 |
| Clostridium_sporogenes_66_CBOT_289 | I | 250 | 4220556 | 35618 | 36 |
| Clostridium_sporogenes_8_O_428 | I | 19 | 4088089 | 929532 | 2 |
| Clostridium_sporogenes_87_0535_260 | I | 202 | 4021062 | 46656 | 27 |
| Clostridium_sporogenes_88_0163_261 | I | 213 | 4101907 | 40700 | 31 |
| Clostridium_sporogenes_ATCC_15579_30 | I | 2 | 4102325 | 2710723 | 1 |
| Clostridium_sporogenes_ATCC_19404_396 | I | 48 | 4066128 | 344087 | 5 |
| Clostridium_sporogenes_DSM_795_283 | I | 1 | 4142990 | 4142990 | 1 |
| Clostridium_sporogenes_NCIMB_10696_279 | I | 1 | 4141984 | 4141984 | 1 |
| Clostridium_sporogenes_PA_3679_1961_4_346 | I | 85 | 3968020 | 115686 | 11 |
| Clostridium_sporogenes_PA_3679_67 | I | 107 | 4180273 | 172248 | 8 |
| Clostridium_sporogenes_PA_3679_Camp_350 | I | 57 | 3984395 | 159201 | 9 |
| Clostridium_sporogenes_PA_3679_FDA_348 | I | 86 | 4021963 | 151708 | 10 |
| Clostridium_sporogenes_PA_3679_NFL_349 | I | 79 | 3959602 | 136601 | 12 |
| Clostridium_sporogenes_PA_3679_UW_347 | I | 48 | 3926962 | 150182 | 10 |
| Clostridium_sporogenes_UC9000_324 | I | 111 | 4338649 | 110240 | 13 |
| Clostridium_tepidiprofundi_DSM_19306_427 | I | 175 | 3060113 | 65328 | 13 |
| Clostridium_tetani_184.08_235 | I | 152 | 2914338 | 70366 | 14 |
| Clostridium_tetani_3911_TET3911 | I | 107 | 2874950 | 263604 | 4 |
| Clostridium_tetani_A_243 | I | 93 | 2824295 | 96331 | 10 |
| Clostridium_tetani_ATCC_19406_217 | I | 50 | 2789793 | 181872 | 5 |
| Clostridium_tetani_ATCC_453_219 | I | 40 | 2890535 | 253841 | 6 |
| Clostridium_tetani_ATCC_454_281 | I | 67 | 2852621 | 87223 | 11 |
| Clostridium_tetani_ATCC_9441_218 | I | 28 | 2800144 | 1734923 | 1 |
| Clostridium_tetani_C2_220 | I | 35 | 2829019 | 263508 | 4 |
| Clostridium_tetani_CN655_234 | I | 118 | 2850319 | 110001 | 9 |
| Clostridium_tetani_E88_Massachusetts | I | 2 | 2873333 | 2799251 | 1 |
| Clostridium_tunisiense_TJ_78 | I | 60 | 4308385 | 341305 | 5 |
| Clostridium_tyrobutyricum_DIVETGP_148 | I | 45 | 3018999 | 168670 | 6 |
| Clostridium_tyrobutyricum_DSM_2637_ATCC_25755_JCM_11008_124 | I | 44 | 3022704 | 133576 | 8 |
| Clostridium_tyrobutyricum_FAM22552_238 | I | 58 | 3052855 | 166858 | 7 |
| Clostridium_tyrobutyricum_FAM22553_239 | I | 62 | 3085051 | 187297 | 6 |
| Clostridium_tyrobutyricum_IFP923_359 | I | 139 | 3190249 | 69218 | 14 |
| Clostridium_tyrobutyricum_KCTC_5387_433 | I | 2 | 3134437 | 3071606 | 1 |
| Clostridium_tyrobutyricum_UC7086_85 | I | 110 | 3064215 | 70370 | 14 |
| Clostridium_ventriculi_17_358SARCINA | I | 32 | 2479673 | 260452 | 3 |
| Clostridium_ventriculi_2789STDY5834858_332SARCINA | I | 31 | 2457263 | 342082 | 3 |
| Clostridium_ventriculi_357SARCINA | I | 91 | 2474230 | 74523 | 11 |
| Clostridium_autoethanogenum_DSM_10061_356 | I | 1 | 4352446 | 4352446 | 1 |
| Clostridium_ragsdalei_P11_436 | I | 79 | 4424992 | 119012 | 11 |
| Clostridium_saudiense_JCC_147 | I | 100 | 3653762 | 89852 | 13 |
| Clostridium_senegalense_JC122_type_strain_JC122_76 | I | 83 | 3925888 | 721927 | 2 |
| Clostridium_sp._7_2_43FAA_34 | I | 5 | 3813122 | 3226588 | 1 |
| Clostridium_sp._Ade.TY_135 | I | 66 | 3113901 | 139705 | 7 |
| Clostridium_sp._ATCC_25772_397 | I | 241 | 4009413 | 62088 | 18 |
| Clostridium_sp._BL_8_531 | I | 231 | 6045940 | 56741 | 35 |
| Clostridium_sp._C8_282 | I | 125 | 4020958 | 59667 | 16 |
| Clostridium_sp._CL_6_212 | I | 17 | 4325182 | 695847 | 2 |
| Clostridium_sp._CL_6_human_stool_245 | I | 17 | 4325182 | 695847 | 2 |
| Clostridium_sp._DL_VIII_62 | I | 1 | 6477357 | 6477357 | 1 |
| Clostridium_sp._HMSC19A10_461 | I | 100 | 4538577 | 125680 | 11 |
| Clostridium_sp._IBUN125C_276 | I | 71 | 4596888 | 133994 | 10 |
| Clostridium_sp._IBUN13A_277 | I | 261 | 4643590 | 42668 | 29 |
| Clostridium_sp._IBUN22A_275 | I | 208 | 4607385 | 41410 | 33 |
| Clostridium_sp._IBUN62F_274 | I | 85 | 3836807 | 120226 | 11 |
| Clostridium_sp._L74_321 | I | 81 | 3687211 | 123171 | 9 |
| Clostridium_sp._LF2_214 | I | 15 | 3750216 | 2057217 | 1 |
| Clostridium_sp._Marseille_P2414_534 | I | 4 | 3799489 | 3796548 | 1 |
| Clostridium_sp._Marseille_P2434_536 | I | 5 | 3083338 | 2631890 | 1 |
| Clostridium_sp._ND2_329 | I | 11 | 3732966 | 2114283 | 1 |
| Clostridium_sulfidigenes_113A_207 | I | 96 | 3717420 | 48174 | 26 |
| Clostridium_cellobioparum_DSM_1351_ATCC_15832_158 | III | 80 | 6132222 | 194099 | 12 |
| Clostridium_cellulolyticum_H10_17 | III | 1 | 4068724 | 4068724 | 1 |
| Clostridium_clariflavum_4_2a_141 | III | 4 | 4872398 | 4415514 | 1 |
| Clostridium_clariflavum_DSM_19732_66 | III | 1 | 4897678 | 4897678 | 1 |
| Clostridium_josui_JCM_17888_143 | III | 2 | 4469680 | 3589078 | 1 |
| Clostridium_papyrosolvens_DSM_2782_50 | III | 31 | 4915287 | 279057 | 6 |
| Clostridium_stercorarium_subsp._leptospartum_DSM_9219_442 | III | 1 | 3148357 | 3148357 | 1 |
| Clostridium_stercorarium_subsp._stercorarium_DSM_8532_88 | III | 1 | 2970010 | 2970010 | 1 |
| Clostridium_stercorarium_subsp._thermolacticum_DSM_2910_441 | III | 1 | 3035622 | 2970010 | 1 |
| Clostridium_termitidis_CT1112_89 | III | 78 | 6415858 | 146289 | 15 |
| Clostridium_sp._Bc_iso_3_444 | III | 4 | 4327139 | 4269335 | 1 |
| Clostridium_sp._BNL1100_69 | III | 1 | 4613747 | 4613747 | 1 |
| Clostridium_leptum_DSM_753_25 | iv /XIVa | 21 | 3270209 | 452649 | 4 |
| Clostridium_sporosphaeroides_DSM_1294_VPI_4527_111 | IV/XIVa | 21 | 3174421 | 324437 | 4 |
| Clostridioides_difficile_VL_0092_988 | XIa | 321 | 4157070 | 90393 | 16 |
| Clostridium_difficile_002_P50_2011_574 | XIa | 79 | 4103061 | 207265 | 8 |
| Clostridium_difficile_01A09CD0020_822 | XIa | 103 | 4211492 | 116608 | 13 |
| Clostridium_difficile_050_P50_2011_576 | XIa | 65 | 4227225 | 211060 | 8 |
| Clostridium_difficile_08ACD0030_812 | XIa | 1 | 4167076 | 4167076 | 1 |
| Clostridium_difficile_103_815 | XIa | 162 | 4012303 | 131308 | 9 |
| Clostridium_difficile_106_814 | XIa | 106 | 4044391 | 137376 | 10 |
| Clostridium_difficile_133_816 | XIa | 150 | 4152605 | 137573 | 10 |
| Clostridium_difficile_20100211_824 | XIa | 5 | 4189946 | 2845247 | 1 |
| Clostridium_difficile_20100502_825 | XIa | 4 | 4201508 | 3853484 | 1 |
| Clostridium_difficile_20110270_833 | XIa | 4 | 4179937 | 3881918 | 1 |
| Clostridium_difficile_20110740_834 | XIa | 9 | 4279635 | 3058973 | 1 |
| Clostridium_difficile_20110995_836 | XIa | 2 | 4092490 | 3015771 | 1 |
| Clostridium_difficile_20111003_837 | XIa | 3 | 4200115 | 2285964 | 1 |
| Clostridium_difficile_20111144_831 | XIa | 13 | 4259046 | 2732843 | 1 |
| Clostridium_difficile_20121412_842 | XIa | 4 | 4278990 | 2484966 | 1 |
| Clostridium_difficile_22_1_820 | XIa | 58 | 4180898 | 196766 | 7 |
| Clostridium_difficile_5_3_790 | XIa | 27 | 4009318 | 786725 | 2 |
| Clostridium_difficile_6042_636 | XIa | 314 | 3945922 | 24692 | 46 |
| Clostridium_difficile_630_561 | XIa | 2 | 4298133 | 4290252 | 1 |
| Clostridium_difficile_630Derm_800 | XIa | 1 | 4293049 | 4293049 | 1 |
| Clostridium_difficile_655_631 | XIa | 207 | 4081576 | 36927 | 35 |
| Clostridium_difficile_7_10492_818 | XIa | 74 | 4317574 | 215323 | 7 |
| Clostridium_difficile_848 | XIa | 159 | 4051221 | 96828 | 14 |
| Clostridium_difficile_BR81_843 | XIa | 1 | 4124384 | 4124384 | 1 |
| Clostridium_difficile_CD03_1454 | XIa | 53 | 3841898 | 147309 | 9 |
| Clostridium_difficile_CD04_1396 | XIa | 61 | 4254923 | 170309 | 9 |
| Clostridium_difficile_CD05_1411 | XIa | 34 | 4443217 | 277295 | 4 |
| Clostridium_difficile_CD06_1403 | XIa | 42 | 4127790 | 231706 | 6 |
| Clostridium_difficile_CD07_1422 | XIa | 32 | 4132581 | 246756 | 7 |
| Clostridium_difficile_CD08_1449 | XIa | 53 | 4224446 | 199231 | 7 |
| Clostridium_difficile_CD10_165_808 | XIa | 67 | 4277571 | 210313 | 7 |
| Clostridium_difficile_CD105HE1_849 | XIa | 46 | 4153465 | 230007 | 7 |
| Clostridium_difficile_CD105KSE2_1198 | XIa | 122 | 4228132 | 180336 | 9 |
| Clostridium_difficile_CD105KSE3_1199 | XIa | 189 | 4295063 | 161100 | 9 |
| Clostridium_difficile_CD105KSE5_1197 | XIa | 378 | 4395161 | 277022 | 6 |
| Clostridium_difficile_CD105KSE6_1196 | XIa | 124 | 4262632 | 190962 | 7 |
| Clostridium_difficile_CD105KSO8_1201 | XIa | 90 | 4521395 | 221277 | 6 |
| Clostridium_difficile_CD111_700 | XIa | 240 | 4047257 | 35117 | 38 |
| Clostridium_difficile_CD113_701 | XIa | 127 | 4255649 | 82434 | 18 |
| Clostridium_difficile_CD12_1446 | XIa | 23 | 4076947 | 821395 | 2 |
| Clostridium_difficile_CD13_1445 | XIa | 39 | 4132205 | 229911 | 6 |
| Clostridium_difficile_CD131_608 | XIa | 134 | 4320519 | 57300 | 27 |
| Clostridium_difficile_CD144_611 | XIa | 56 | 4095302 | 184803 | 8 |
| Clostridium_difficile_CD15_1452 | XIa | 52 | 4155465 | 209526 | 6 |
| Clostridium_difficile_CD159_613 | XIa | 290 | 3986487 | 24265 | 52 |
| Clostridium_difficile_CD166_616 | XIa | 56 | 4289366 | 222760 | 8 |
| Clostridium_difficile_CD18_1435 | XIa | 39 | 4142242 | 246977 | 5 |
| Clostridium_difficile_CD19_1458 | XIa | 62 | 4022263 | 140793 | 10 |
| Clostridium_difficile_CD196_562 | XIa | 1 | 4110554 | 4110554 | 1 |
| Clostridium_difficile_CD21_1414 | XIa | 43 | 4125071 | 229812 | 6 |
| Clostridium_difficile_CD211_628 | XIa | 187 | 4054786 | 42827 | 30 |
| Clostridium_difficile_CD212_629 | XIa | 36 | 4003929 | 240914 | 6 |
| Clostridium_difficile_CD22_1423 | XIa | 35 | 4178020 | 247845 | 5 |
| Clostridium_difficile_CD23_1419 | XIa | 38 | 4061847 | 263841 | 5 |
| Clostridium_difficile_CD24_1407 | XIa | 44 | 4168771 | 204915 | 7 |
| Clostridium_difficile_CD25_1412 | XIa | 41 | 4293106 | 228970 | 7 |
| Clostridium_difficile_CD26_1388 | XIa | 26 | 4014727 | 624690 | 3 |
| Clostridium_difficile_CD26A54_R_809 | XIa | 4 | 4146282 | 1881735 | 2 |
| Clostridium_difficile_CD26A54_S_810 | XIa | 6 | 4166728 | 1445886 | 2 |
| Clostridium_difficile_CD27_1391 | XIa | 41 | 4197119 | 265736 | 7 |
| Clostridium_difficile_CD28_1392 | XIa | 54 | 4157621 | 201745 | 7 |
| Clostridium_difficile_CD30_1400 | XIa | 47 | 4124166 | 188065 | 9 |
| Clostridium_difficile_CD31_1437 | XIa | 34 | 4236825 | 283093 | 5 |
| Clostridium_difficile_CD35_1438 | XIa | 39 | 4025541 | 264934 | 4 |
| Clostridium_difficile_CD39_1432 | XIa | 30 | 4251978 | 402273 | 3 |
| Clostridium_difficile_CD41_1448 | XIa | 30 | 4181594 | 438667 | 4 |
| Clostridium_difficile_CD42_1416 | XIa | 28 | 4307274 | 476381 | 3 |
| Clostridium_difficile_CD43_1428 | XIa | 54 | 3909083 | 141853 | 10 |
| Clostridium_difficile_CD44_1431 | XIa | 49 | 4429426 | 216355 | 6 |
| Clostridium_difficile_CD45_597 | XIa | 176 | 4367906 | 54395 | 26 |
| Clostridium_difficile_CD46_1447 | XIa | 24 | 4077061 | 821490 | 2 |
| Clostridium_difficile_CD49_1401 | XIa | 59 | 4208917 | 245737 | 6 |
| Clostridium_difficile_CD51_1417 | XIa | 41 | 4215495 | 274737 | 6 |
| Clostridium_difficile_CD52_1420 | XIa | 47 | 4417523 | 212914 | 6 |
| Clostridium_difficile_CD53_1390 | XIa | 25 | 4079056 | 823366 | 2 |
| Clostridium_difficile_CD54_1402 | XIa | 34 | 4214944 | 633620 | 3 |
| Clostridium_difficile_CD57_1444 | XIa | 34 | 4313723 | 300939 | 4 |
| Clostridium_difficile_CD58_1427 | XIa | 32 | 4196484 | 383688 | 4 |
| Clostridium_difficile_CD60_1399 | XIa | 54 | 4122619 | 192092 | 9 |
| Clostridium_difficile_CD62_1397 | XIa | 27 | 4067368 | 405787 | 3 |
| Clostridium_difficile_CD63_1387 | XIa | 32 | 4012749 | 262729 | 4 |
| Clostridium_difficile_CD64_1425 | XIa | 38 | 4154464 | 268013 | 6 |
| Clostridium_difficile_CD65_1393 | XIa | 39 | 4101999 | 247554 | 6 |
| Clostridium_difficile_CD66_1406 | XIa | 28 | 4068751 | 406103 | 3 |
| Clostridium_difficile_CD67_1405 | XIa | 55 | 3961761 | 153556 | 8 |
| Clostridium_difficile_CD68_602 | XIa | 141 | 3975975 | 53580 | 22 |
| Clostridium_difficile_CD69_1395 | XIa | 27 | 4079436 | 357688 | 3 |
| Clostridium_difficile_CD70_1451 | XIa | 33 | 4074098 | 230885 | 6 |
| Clostridium_difficile_CD8_15_813 | XIa | 44 | 4249791 | 235991 | 5 |
| Clostridium_difficile_CD8_586 | XIa | 280 | 4044267 | 28010 | 48 |
| Clostridium_difficile_CD9_587 | XIa | 87 | 4310458 | 685892 | 3 |
| Clostridium_difficile_CD90_697 | XIa | 123 | 4029007 | 74899 | 16 |
| Clostridium_difficile_DA00114_641 | XIa | 165 | 4058203 | 46799 | 27 |
| Clostridium_difficile_DA00126_642 | XIa | 91 | 4167768 | 115337 | 12 |
| Clostridium_difficile_DA00128_643 | XIa | 95 | 4217009 | 88258 | 15 |
| Clostridium_difficile_DA00132_646 | XIa | 78 | 4070905 | 99187 | 14 |
| Clostridium_difficile_DA00134_647 | XIa | 98 | 4115079 | 107219 | 11 |
| Clostridium_difficile_DA00141_648 | XIa | 52 | 4089728 | 130697 | 10 |
| Clostridium_difficile_DA00142_649 | XIa | 193 | 4183071 | 44831 | 33 |
| Clostridium_difficile_DA00145_650 | XIa | 132 | 4051066 | 55379 | 25 |
| Clostridium_difficile_DA00154_652 | XIa | 102 | 4126663 | 109889 | 14 |
| Clostridium_difficile_DA00160_653 | XIa | 98 | 4134851 | 113054 | 13 |
| Clostridium_difficile_DA00183_656 | XIa | 114 | 4071468 | 67760 | 20 |
| Clostridium_difficile_DA00193_658 | XIa | 60 | 3969329 | 141166 | 9 |
| Clostridium_difficile_DA00195_659 | XIa | 95 | 4108697 | 95853 | 16 |
| Clostridium_difficile_DA00196_660 | XIa | 22 | 4254880 | 623272 | 3 |
| Clostridium_difficile_DA00197_661 | XIa | 18 | 4122528 | 451615 | 4 |
| Clostridium_difficile_DA00211_664 | XIa | 90 | 4072757 | 88029 | 16 |
| Clostridium_difficile_DA00238_669 | XIa | 94 | 4163916 | 96856 | 14 |
| Clostridium_difficile_DA00244_670 | XIa | 15 | 4067645 | 761856 | 2 |
| Clostridium_difficile_DA00245_671 | XIa | 336 | 3985935 | 20188 | 61 |
| Clostridium_difficile_DA00273_607 | XIa | 271 | 4095984 | 25512 | 47 |
| Clostridium_difficile_DA00307_675 | XIa | 219 | 4147275 | 36095 | 37 |
| Clostridium_difficile_DA00313_677 | XIa | 112 | 4083557 | 70240 | 20 |
| Clostridium_difficile_E1_789 | XIa | 212 | 3945725 | 38456 | 31 |
| Clostridium_difficile_E12_782 | XIa | 353 | 4009421 | 22652 | 55 |
| Clostridium_difficile_E13_773 | XIa | 262 | 4146240 | 41362 | 31 |
| Clostridium_difficile_E14_767 | XIa | 319 | 4131267 | 32211 | 40 |
| Clostridium_difficile_E23_766 | XIa | 264 | 4015392 | 37968 | 34 |
| Clostridium_difficile_E9_786 | XIa | 373 | 4156306 | 29903 | 41 |
| Clostridium_difficile_F480_748 | XIa | 32 | 3989604 | 241744 | 6 |
| Clostridium_difficile_F548_750 | XIa | 17 | 4257706 | 695337 | 3 |
| Clostridium_difficile_F601_751 | XIa | 40 | 4103199 | 214607 | 8 |
| Clostridium_difficile_G46_794 | XIa | 1 | 4189317 | 4189317 | 1 |
| Clostridium_difficile_H3_819 | XIa | 76 | 4138192 | 217879 | 7 |
| Clostridium_difficile_KY62_802 | XIa | 277 | 4040078 | 187206 | 7 |
| Clostridium_difficile_KY64_803 | XIa | 324 | 4035637 | 180982 | 7 |
| Clostridium_difficile_LIBA_5784_797 | XIa | 44 | 4156825 | 234855 | 6 |
| Clostridium_difficile_M68_572 | XIa | 1 | 4308325 | 4308325 | 1 |
| Clostridium_difficile_NAP07_571 | XIa | 33 | 3905466 | 525841 | 4 |
| Clostridium_difficile_P1_687 | XIa | 160 | 4104507 | 48238 | 28 |
| Clostridium_difficile_P21_674 | XIa | 53 | 4090537 | 162302 | 9 |
| Clostridium_difficile_P32_726 | XIa | 44 | 4027255 | 237219 | 7 |
| Clostridium_difficile_P33_755 | XIa | 57 | 4103431 | 150350 | 9 |
| Clostridium_difficile_P36_728 | XIa | 40 | 4048210 | 233587 | 8 |
| Clostridium_difficile_P37_757 | XIa | 62 | 4080614 | 180173 | 8 |
| Clostridium_difficile_P41_756 | XIa | 62 | 4119865 | 234033 | 6 |
| Clostridium_difficile_P42_729 | XIa | 53 | 4185941 | 195731 | 7 |
| Clostridium_difficile_P46_731 | XIa | 60 | 4125436 | 196884 | 9 |
| Clostridium_difficile_P48_732 | XIa | 68 | 4172979 | 157205 | 9 |
| Clostridium_difficile_P49_733 | XIa | 46 | 4153887 | 235599 | 8 |
| Clostridium_difficile_P5_689 | XIa | 181 | 4048946 | 44667 | 32 |
| Clostridium_difficile_P50_734 | XIa | 29 | 4395649 | 486408 | 4 |
| Clostridium_difficile_P51_735 | XIa | 56 | 4256776 | 161476 | 10 |
| Clostridium_difficile_P53_758 | XIa | 41 | 4025043 | 236448 | 8 |
| Clostridium_difficile_P6_673 | XIa | 167 | 4167114 | 48839 | 28 |
| Clostridium_difficile_P64_759 | XIa | 72 | 4226724 | 184824 | 8 |
| Clostridium_difficile_P68_760 | XIa | 49 | 4056190 | 246498 | 7 |
| Clostridium_difficile_P69_739 | XIa | 49 | 4213133 | 228563 | 8 |
| Clostridium_difficile_P70_740 | XIa | 48 | 4049371 | 193767 | 8 |
| Clostridium_difficile_P71_742 | XIa | 52 | 4130145 | 163133 | 8 |
| Clostridium_difficile_P72_741 | XIa | 44 | 4057420 | 207373 | 7 |
| Clostridium_difficile_P75_745 | XIa | 54 | 4050593 | 191288 | 9 |
| Clostridium_difficile_P77_746 | XIa | 61 | 4155874 | 188876 | 8 |
| Clostridium_difficile_P78_738 | XIa | 60 | 4119601 | 181575 | 8 |
| Clostridium_difficile_P8_715 | XIa | 203 | 4292666 | 44870 | 28 |
| Clostridium_difficile_QCD_37x79_QCD_37x79_566 | XIa | 15 | 4332988 | 4092698 | 1 |
| Clostridium_difficile_QCD_63q42_QCD_63q42_563 | XIa | 28 | 4443737 | 4129288 | 1 |
| Clostridium_difficile_QCD_76w55_QCD_76w55_564 | XIa | 24 | 4396895 | 4078976 | 1 |
| Clostridium_difficile_QCD_97b34_QCD_97b34_565 | XIa | 15 | 4063610 | 3998408 | 1 |
| Clostridium_difficile_RA09_70_806 | XIa | 116 | 4232226 | 105769 | 13 |
| Clostridium_difficile_SA10_050_807 | XIa | 81 | 4319873 | 128043 | 11 |
| Clostridium_difficile_SG12_P1_791 | XIa | 102 | 4268551 | 146259 | 11 |
| Clostridium_difficile_T20_762 | XIa | 210 | 3815975 | 38353 | 30 |
| Clostridium_difficile_T22_771 | XIa | 332 | 4083325 | 32040 | 43 |
| Clostridium_difficile_T23_770 | XIa | 293 | 4078839 | 30654 | 37 |
| Clostridium_difficile_T3_776 | XIa | 275 | 4060583 | 34614 | 37 |
| Clostridium_difficile_T61_787 | XIa | 82 | 4071443 | 141610 | 9 |
| Clostridium_difficile_VL_0042_1181 | XIa | 377 | 4132263 | 66649 | 21 |
| Clostridium_difficile_VL_0048_974 | XIa | 245 | 4175645 | 199899 | 8 |
| Clostridium_difficile_VL_0052_907 | XIa | 214 | 4208030 | 161786 | 9 |
| Clostridium_difficile_VL_0059_1016 | XIa | 322 | 3880153 | 67831 | 18 |
| Clostridium_difficile_VL_0088_931 | XIa | 317 | 4456714 | 134792 | 10 |
| Clostridium_difficile_VL_0094_873 | XIa | 299 | 4119551 | 67547 | 18 |
| Clostridium_difficile_VL_0095_989 | XIa | 261 | 4197961 | 142480 | 9 |
| Clostridium_difficile_VL_0104_1105 | XIa | 261 | 4068388 | 88166 | 14 |
| Clostridium_difficile_VL_0108_993 | XIa | 322 | 4117060 | 106650 | 14 |
| Clostridium_difficile_VL_0114_995 | XIa | 339 | 4114836 | 59007 | 23 |
| Clostridium_difficile_VL_0119_998 | XIa | 340 | 4124686 | 83417 | 18 |
| Clostridium_difficile_VL_0123_1000 | XIa | 267 | 4394265 | 65991 | 20 |
| Clostridium_difficile_VL_0135_917 | XIa | 218 | 4282195 | 202085 | 8 |
| Clostridium_difficile_VL_0174_1012 | XIa | 266 | 4156515 | 134544 | 9 |
| Clostridium_difficile_VL_0177_947 | XIa | 306 | 4124242 | 95020 | 16 |
| Clostridium_difficile_VL_0199_1035 | XIa | 266 | 4120144 | 108096 | 13 |
| Clostridium_difficile_VL_0232_1051 | XIa | 213 | 4123326 | 166166 | 10 |
| Clostridium_difficile_VL_0245_1065 | XIa | 315 | 4176184 | 90332 | 15 |
| Clostridium_difficile_VL_0259_941 | XIa | 376 | 4098661 | 52866 | 22 |
| Clostridium_difficile_VL_0291_896 | XIa | 278 | 3881849 | 90247 | 16 |
| Clostridium_difficile_VL_0305_1080 | XIa | 310 | 4159239 | 149863 | 8 |
| Clostridium_difficile_VL_0307_1081 | XIa | 255 | 4123594 | 195946 | 6 |
| Clostridium_difficile_VL_0308_1082 | XIa | 280 | 4239945 | 71640 | 18 |
| Clostridium_difficile_VL_0311_884 | XIa | 388 | 4254985 | 44720 | 29 |
| Clostridium_difficile_VL_0359_1101 | XIa | 215 | 4095701 | 58622 | 19 |
| Clostridium_difficile_VL_0404_1130 | XIa | 233 | 4088651 | 219919 | 7 |
| Clostridium_difficile_VL_0414_1134 | XIa | 369 | 4124988 | 56163 | 22 |
| Clostridium_difficile_VL_0417_1149 | XIa | 343 | 4130169 | 129218 | 10 |
| Clostridium_difficile_VL_0426_1188 | XIa | 370 | 4204351 | 70763 | 15 |
| Clostridium_difficile_VL_0429_891 | XIa | 323 | 4113210 | 46838 | 25 |
| Clostridium_difficile_VL_0452_1104 | XIa | 293 | 4408841 | 135522 | 12 |
| Clostridium_difficile_VL_0459_944 | XIa | 344 | 4382319 | 65636 | 21 |
| Clostridium_difficile_VL_0460_1139 | XIa | 325 | 4130158 | 114096 | 11 |
| Clostridium_difficile_VRECD0001_1264 | XIa | 48 | 4255504 | 799464 | 2 |
| Clostridium_difficile_VRECD0003_1255 | XIa | 52 | 4105252 | 219704 | 5 |
| Clostridium_difficile_VRECD0004_1262 | XIa | 25 | 4036206 | 652602 | 3 |
| Clostridium_difficile_VRECD0005_1265 | XIa | 27 | 4256197 | 675134 | 3 |
| Clostridium_difficile_VRECD0006_1268 | XIa | 28 | 4203969 | 598150 | 3 |
| Clostridium_difficile_VRECD0007_1267 | XIa | 52 | 4569562 | 635337 | 3 |
| Clostridium_difficile_VRECD0008_1263 | XIa | 32 | 4093125 | 414552 | 3 |
| Clostridium_difficile_VRECD0009_1257 | XIa | 54 | 4317067 | 473305 | 3 |
| Clostridium_difficile_VRECD0010_1233 | XIa | 27 | 4130133 | 592811 | 3 |
| Clostridium_difficile_VRECD0012_1258 | XIa | 22 | 4126592 | 678324 | 3 |
| Clostridium_difficile_VRECD0014_1260 | XIa | 26 | 4070644 | 567836 | 3 |
| Clostridium_difficile_VRECD0016_1261 | XIa | 31 | 4241280 | 347042 | 4 |
| Clostridium_difficile_VRECD0017_1350 | XIa | 27 | 4149640 | 535845 | 3 |
| Clostridium_difficile_VRECD0019_1344 | XIa | 27 | 4146662 | 306509 | 4 |
| Clostridium_difficile_VRECD0020_1346 | XIa | 33 | 4240856 | 498190 | 3 |
| Clostridium_difficile_VRECD0021_1345 | XIa | 25 | 4011448 | 613070 | 3 |
| Clostridium_difficile_VRECD0023_1347 | XIa | 47 | 4593015 | 301073 | 3 |
| Clostridium_difficile_VRECD0024_1349 | XIa | 26 | 4140268 | 599831 | 3 |
| Clostridium_difficile_VRECD0025_1357 | XIa | 27 | 4161988 | 686847 | 2 |
| Clostridium_difficile_VRECD0026_1348 | XIa | 32 | 4421543 | 599187 | 3 |
| Clostridium_difficile_VRECD0027_1355 | XIa | 30 | 4297925 | 380836 | 5 |
| Clostridium_difficile_VRECD0030_1359 | XIa | 28 | 4512569 | 543500 | 3 |
| Clostridium_difficile_VRECD0033_1361 | XIa | 29 | 4523828 | 593675 | 2 |
| Clostridium_difficile_VRECD0034_1351 | XIa | 27 | 4084869 | 402072 | 3 |
| Clostridium_difficile_VRECD0035_1354 | XIa | 40 | 4233861 | 556145 | 3 |
| Clostridium_difficile_VRECD0036_1360 | XIa | 31 | 4079714 | 588526 | 3 |
| Clostridium_difficile_VRECD0038_1271 | XIa | 23 | 4121922 | 683887 | 2 |
| Clostridium_difficile_VRECD0039_1237 | XIa | 39 | 4249962 | 321186 | 3 |
| Clostridium_difficile_VRECD0041_1273 | XIa | 39 | 4357659 | 421412 | 4 |
| Clostridium_difficile_VRECD0042_1274 | XIa | 38 | 4169930 | 385482 | 4 |
| Clostridium_difficile_VRECD0043_1206 | XIa | 27 | 4139048 | 539782 | 2 |
| Clostridium_difficile_VRECD0047_1276 | XIa | 29 | 4154535 | 475562 | 4 |
| Clostridium_difficile_VRECD0048_1277 | XIa | 26 | 4065912 | 562478 | 3 |
| Clostridium_difficile_VRECD0049_1278 | XIa | 36 | 4205910 | 579078 | 3 |
| Clostridium_difficile_VRECD0051_1279 | XIa | 42 | 4198487 | 402206 | 4 |
| Clostridium_difficile_VRECD0053_1281 | XIa | 30 | 4308716 | 560877 | 2 |
| Clostridium_difficile_VRECD0054_1284 | XIa | 49 | 4209694 | 402151 | 4 |
| Clostridium_difficile_VRECD0055_1283 | XIa | 30 | 4196308 | 693711 | 2 |
| Clostridium_difficile_VRECD0057_1363 | XIa | 23 | 4121104 | 580367 | 3 |
| Clostridium_difficile_VRECD0058_1364 | XIa | 30 | 4191407 | 497568 | 3 |
| Clostridium_difficile_VRECD0059_1369 | XIa | 32 | 4187721 | 562268 | 3 |
| Clostridium_difficile_VRECD0060_1365 | XIa | 36 | 4189636 | 314543 | 5 |
| Clostridium_difficile_VRECD0061_1370 | XIa | 29 | 4121024 | 542064 | 4 |
| Clostridium_difficile_VRECD0063_1322 | XIa | 30 | 4185963 | 561804 | 3 |
| Clostridium_difficile_VRECD0064_1372 | XIa | 26 | 4184952 | 561836 | 3 |
| Clostridium_difficile_VRECD0065_1373 | XIa | 27 | 4187935 | 562286 | 3 |
| Clostridium_difficile_VRECD0067_1367 | XIa | 33 | 4194047 | 687659 | 3 |
| Clostridium_difficile_VRECD0070_1374 | XIa | 36 | 4239257 | 311198 | 4 |
| Clostridium_difficile_VRECD0071_1375 | XIa | 42 | 4237804 | 511935 | 3 |
| Clostridium_difficile_VRECD0072_1376 | XIa | 30 | 4186963 | 562426 | 3 |
| Clostridium_difficile_VRECD0073_1286 | XIa | 33 | 4320443 | 568445 | 3 |
| Clostridium_difficile_VRECD0078_1243 | XIa | 45 | 3981831 | 187468 | 7 |
| Clostridium_difficile_VRECD0082_1250 | XIa | 41 | 4308862 | 239429 | 5 |
| Clostridium_difficile_VRECD0083_1291 | XIa | 43 | 4387145 | 262957 | 5 |
| Clostridium_difficile_VRECD0084_1293 | XIa | 46 | 4389239 | 500411 | 3 |
| Clostridium_difficile_VRECD0085_1251 | XIa | 30 | 4074943 | 380392 | 4 |
| Clostridium_difficile_VRECD0089_1324 | XIa | 33 | 4114466 | 296534 | 6 |
| Clostridium_difficile_VRECD0095_1326 | XIa | 28 | 4174911 | 257511 | 6 |
| Clostridium_difficile_VRECD0099_1295 | XIa | 29 | 4165342 | 483795 | 4 |
| Clostridium_difficile_VRECD0100_1294 | XIa | 34 | 4419253 | 600146 | 3 |
| Clostridium_difficile_VRECD0101_1296 | XIa | 28 | 4274243 | 485062 | 4 |
| Clostridium_difficile_VRECD0102_1297 | XIa | 31 | 4098928 | 597153 | 3 |
| Clostridium_difficile_VRECD0103_1298 | XIa | 32 | 4199524 | 508865 | 3 |
| Clostridium_difficile_VRECD0106_1301 | XIa | 23 | 4045108 | 653419 | 2 |
| Clostridium_difficile_VRECD0109_1385 | XIa | 30 | 4098893 | 563074 | 3 |
| Clostridium_difficile_VRECD0110_1386 | XIa | 50 | 4238718 | 498953 | 3 |
| Clostridium_difficile_VRECD0112_1302 | XIa | 40 | 4266439 | 561758 | 3 |
| Clostridium_difficile_VRECD0115_1305 | XIa | 28 | 4096161 | 546233 | 3 |
| Clostridium_difficile_VRECD0116_1327 | XIa | 97 | 4571183 | 167119 | 11 |
| Clostridium_difficile_VRECD0119_1229 | XIa | 41 | 4347268 | 789394 | 3 |
| Clostridium_difficile_VRECD0122_1252 | XIa | 47 | 4114017 | 179488 | 7 |
| Clostridium_difficile_VRECD0123_1223 | XIa | 38 | 3945589 | 202201 | 5 |
| Clostridium_difficile_VRECD0127_1230 | XIa | 40 | 4431161 | 707933 | 3 |
| Clostridium_difficile_VRECD0128_1238 | XIa | 357 | 4448160 | 598450 | 3 |
| Clostridium_difficile_VRECD0138_1213 | XIa | 37 | 4291619 | 362902 | 5 |
| Clostridium_difficile_VRECD0139_1236 | XIa | 35 | 4446093 | 682760 | 3 |
| Clostridium_difficile_VRECD0140_1224 | XIa | 40 | 4282826 | 368740 | 5 |
| Clostridium_difficile_VRECD0141_1211 | XIa | 33 | 4291501 | 450759 | 4 |
| Clostridium_difficile_VRECD0144_1309 | XIa | 35 | 4120089 | 402297 | 4 |
| Clostridium_difficile_VRECD0145_1310 | XIa | 40 | 4210086 | 698250 | 2 |
| Clostridium_difficile_VRECD0146_1311 | XIa | 36 | 4294414 | 526365 | 4 |
| Clostridium_difficile_VRECD0148_1313 | XIa | 65 | 4252825 | 160557 | 10 |
| Clostridium_difficile_VRECD0149_1314 | XIa | 39 | 4116743 | 221331 | 7 |
| Clostridium_difficile_VRECD0153_1330 | XIa | 55 | 4225485 | 343945 | 4 |
| Clostridium_difficile_VRECD0154_1318 | XIa | 38 | 4147832 | 414616 | 3 |
| Clostridium_difficile_VRECD0157_1321 | XIa | 54 | 4397606 | 268671 | 6 |
| Clostridium_difficile_VRECD0158_1329 | XIa | 56 | 4396137 | 205249 | 8 |
| Clostridium_difficile_VRECD0159_1216 | XIa | 34 | 4316578 | 394724 | 4 |
| Clostridium_difficile_VRECD0160_1221 | XIa | 29 | 4178309 | 529907 | 4 |
| Clostridium_difficile_VRECD0162_1226 | XIa | 31 | 4062673 | 308123 | 5 |
| Clostridium_difficile_VRECD0163_1219 | XIa | 37 | 4231218 | 346917 | 4 |
| Clostridium_difficile_VRECD0165_1212 | XIa | 44 | 4248821 | 422442 | 3 |
| Clostridium_difficile_VRECD0166_1249 | XIa | 33 | 4168589 | 505747 | 3 |
| Clostridium_difficile_VRECD0167_1234 | XIa | 55 | 4289170 | 501840 | 4 |
| Clostridium_difficile_VRECD0169_1245 | XIa | 30 | 4057998 | 387103 | 4 |
| Clostridium_difficile_VRECD0170_1244 | XIa | 39 | 4222769 | 477083 | 4 |
| Clostridium_difficile_VRECD0171_1253 | XIa | 38 | 4325883 | 263908 | 7 |
| Clostridium_difficile_VRECD0175_1235 | XIa | 26 | 4366076 | 632922 | 3 |
| Clostridium_difficile_VRECD0177_1222 | XIa | 28 | 4282513 | 454161 | 3 |
| Clostridium_difficile_VRECD0178_1241 | XIa | 37 | 4257551 | 571332 | 4 |
| Clostridium_difficile_VRECD0179_1256 | XIa | 34 | 4338827 | 428504 | 4 |
| Clostridium_difficile_VRECD0180_1332 | XIa | 43 | 4290419 | 222418 | 7 |
| Clostridium_difficile_VRECD0182_1334 | XIa | 37 | 4294031 | 242766 | 7 |
| Clostridium_difficile_VRECD0183_1335 | XIa | 38 | 4173270 | 519461 | 3 |
| Clostridium_difficile_VRECD0184_1336 | XIa | 33 | 4110442 | 512879 | 3 |
| Clostridium_difficile_VRECD0185_1337 | XIa | 30 | 4250524 | 341755 | 4 |
| Clostridium_difficile_VRECD0189_1339 | XIa | 45 | 4245033 | 193494 | 7 |
| Clostridium_difficile_VRECD0190_1338 | XIa | 30 | 4169585 | 271547 | 5 |
| Clostridium_difficile_VRECD0191_1340 | XIa | 48 | 4172317 | 184561 | 9 |
| Clostridium_difficile_Y10_682 | XIa | 37 | 4062917 | 237821 | 6 |
| Clostridium_difficile_Y155_685 | XIa | 232 | 3980239 | 33044 | 36 |
| Clostridium_difficile_Y165_704 | XIa | 367 | 4263767 | 24471 | 55 |
| Clostridium_difficile_Y231_707 | XIa | 42 | 4093690 | 197543 | 8 |
| Clostridium_difficile_Y247_694 | XIa | 44 | 4053085 | 199386 | 7 |
| Clostridium_difficile_Y270_705 | XIa | 53 | 4149527 | 217760 | 8 |
| Clostridium_difficile_Y312_706 | XIa | 160 | 4054373 | 47451 | 26 |
| Clostridium_difficile_Y381_713 | XIa | 60 | 4060729 | 234466 | 8 |
| Clostridium_difficile_Y401_686 | XIa | 205 | 4000151 | 35327 | 36 |
| Clostridium_difficile_Y41_684 | XIa | 39 | 4074379 | 233399 | 7 |
| Clostridium_sp._HMC19D07_459 | Xia | 98 | 4217734 | 139497 | 12 |
| Clostridium_sp._HMSC19A11_449 | Xia | 117 | 4235516 | 136274 | 9 |
| Clostridium_sp._HMSC19B01_448 | Xia | 101 | 4083546 | 118969 | 12 |
| Clostridium_sp._HMSC19B04_451 | Xia | 68 | 3979326 | 184637 | 9 |
| Clostridium_sp._HMSC19B10_450 | Xia | 77 | 4056622 | 128616 | 11 |
| Clostridium_sp._HMSC19B11_452 | Xia | 136 | 4137486 | 91682 | 15 |
| Clostridium_sp._HMSC19B12_453 | Xia | 113 | 4286468 | 138393 | 11 |
| Clostridium_sp._HMSC19C05_454 | Xia | 69 | 4062786 | 118779 | 12 |
| Clostridium_sp._HMSC19C08_455 | Xia | 79 | 4069234 | 128619 | 11 |
| Clostridium_sp._HMSC19C09_456 | Xia | 91 | 4060413 | 113889 | 14 |
| Clostridium_sp._HMSC19C11_457 | Xia | 79 | 4015638 | 152320 | 10 |
| Clostridium_sp._HMSC19D02_458 | Xia | 85 | 4070187 | 157295 | 9 |
| Clostridium_sp._HMSC19E03_460 | Xia | 64 | 4059828 | 134413 | 10 |
| Clostridium_aerotolerans_DSM_5434_165 | XIVa | 50 | 4732373 | 213027 | 7 |
| Clostridium_aminophilum_542 | XIVa | 31 | 3113820 | 176596 | 6 |
| Clostridium_aminophilum_DSM_10710_178 | XIVa | 36 | 3114416 | 176585 | 6 |
| Clostridium_aminophilum_KH1P1_540 | XIVa | 108 | 3198475 | 54860 | 19 |
| Clostridium_bolteae_90A5_110 | XIVa | 12 | 6421395 | 1978325 | 2 |
| Clostridium_bolteae_90A9_109 | XIVa | 1 | 6377378 | 6377378 | 1 |
| Clostridium_bolteae_90B3_108 | XIVa | 4 | 6538460 | 3794250 | 1 |
| Clostridium_bolteae_90B7_107 | XIVa | 19 | 6439235 | 799384 | 4 |
| Clostridium_bolteae_90B8_106 | XIVa | 21 | 6482686 | 663201 | 5 |
| Clostridium_bolteae_ATCC_BAA_613_26 | XIVa | 68 | 6557988 | 187711 | 12 |
| Clostridium_bolteae_WAL_14578_291 | XIVa | 30 | 6604884 | 668224 | 3 |
| Clostridium_celerecrescens_152B_206 | XIVa | 92 | 5038011 | 208566 | 9 |
| Clostridium_citroniae_WAL_17108_63 | XIVa | 41 | 6647380 | 473033 | 3 |
| Clostridium_citroniae_WAL19142_292 | XIVa | 38 | 6252818 | 694783 | 4 |
| Clostridium_clostridioforme_2_1_49FAA_65 | XIVa | 69 | 5500475 | 722831 | 3 |
| Clostridium_clostridioforme_2789STDY5834865_334 | XIVa | 202 | 5514222 | 70965 | 22 |
| Clostridium_clostridioforme_90A1_104 | XIVa | 16 | 5806027 | 1108021 | 2 |
| Clostridium_clostridioforme_90A3_103 | XIVa | 11 | 5549890 | 1495260 | 2 |
| Clostridium_clostridioforme_90A4_102 | XIVa | 48 | 5871489 | 217308 | 10 |
| Clostridium_clostridioforme_90A6_101 | XIVa | 22 | 6033914 | 582564 | 3 |
| Clostridium_clostridioforme_90A7_100 | XIVa | 8 | 6180373 | 1761752 | 2 |
| Clostridium_clostridioforme_90A8_99 | XIVa | 47 | 5974284 | 254783 | 9 |
| Clostridium_clostridioforme_90B1_98 | XIVa | 10 | 5602152 | 1271686 | 2 |
| Clostridium_clostridioforme_AGR2157_122 | XIVa | 133 | 4943165 | 83167 | 20 |
| Clostridium_clostridioforme_ATCC_25537_544 | XIVa | 148 | 5465751 | 80688 | 20 |
| Clostridium_clostridioforme_CM201_95 | XIVa | 7 | 5655915 | 1153171 | 2 |
| Clostridium_clostridioforme_NLAE_zl_C196_539 | XIVa | 164 | 5225716 | 59867 | 27 |
| Clostridium_clostridioforme_NLAE_zl_G208_41 | XIVa | 157 | 5236575 | 65106 | 24 |
| Clostridium_clostridioforme_WAL_7855_293 | XIVa | 70 | 5459495 | 262765 | 5 |
| Clostridium_glycyrrhizinilyticum_JCM_13369_328 | XIVa | 65 | 3215420 | 112462 | 10 |
| Clostridium_hylemonae_DSM_15053_32 | XIVa | 123 | 3889859 | 2898417 | 1 |
| Clostridium_indolis_DSM_755_145 | XIVa | 1 | 6383701 | 6383701 | 1 |
| Clostridium_methoxybenzovorans_SR3_117 | XIVa | 14 | 7085377 | 1607802 | 2 |
| Clostridium_methylpentosum_DSM_5476_35 | XIVa | 17 | 3478423 | 779329 | 2 |
| Clostridium_saccharolyticum_WM1_22 | XIVa | 1 | 4662871 | 4662871 | 1 |
| Clostridium_symbiosum_2789STDY5834864_339 | XIVa | 186 | 4727130 | 102800 | 13 |
| Clostridium_symbiosum_ATCC_14940_129 | XIVa | 270 | 4823675 | 30142 | 49 |
| Clostridium_symbiosum_WAL_14163_54 | XIVa | 52 | 5352498 | 328026 | 6 |
| Clostridium_symbiosum_WAL_14673_55 | XIVa | 55 | 4916964 | 628413 | 4 |
| Clostridium_ihumii_AP5_155 | XIVa | 96 | 4433668 | 124325 | 10 |
| Clostridium_sp._12A_144 | XIVa | 3 | 4605255 | 3532345 | 1 |
| Clostridium_sp._7_3_54FAA_64 | XIVa | 51 | 5464886 | 459666 | 5 |
| Clostridium_sp._ASBs410_154 | XIVa | 1 | 5723672 | 5723672 | 1 |
| Clostridium_sp._ATCC_29733_130 | XIVa | 161 | 3084735 | 45185 | 19 |
| Clostridium_sp._BR31_446 | XIVa | 64 | 3318223 | 154838 | 7 |
| Clostridium_sp._DSM_4029_550 | XIVa | 11 | 3113497 | 984592 | 2 |
| Clostridium_sp._FS41_257 | XIVa | 97 | 6265866 | 227665 | 9 |
| Clostridium_sp._M62_1_36 | XIVa | 26 | 3842594 | 463016 | 3 |
| Clostridium_sp._Marseille_P2415_560 | XIVa | 9 | 5247868 | 4178102 | 1 |
| Clostridium_sp._Marseille_P2538_535 | XIVa | 2 | 4144149 | 4143223 | 1 |
| Clostridium_sp._Marseille_P3244_547 | XIVa | 3 | 2972275 | 2607599 | 1 |
| Clostridium_sp._SY8519_73 | XIVa | 1 | 2835737 | 2835737 | 1 |
| Clostridium_lactatifermentans_DSM_14214_559 | XIVb | 163 | 3546077 | 42744 | 26 |
| Clostridium_neopropionicum_DSM_3847_363 | XIVb | 29 | 3194881 | 311201 | 4 |
| Clostridium_propionicum_DSM_1682_548 | XIVb | 28 | 3076693 | 170419 | 5 |
| Clostridium_propionicum_DSM_1682_X2_362 | XIVb | 1 | 3120417 | 3120417 | 1 |
| Clostridium_sp._ASF356_93 | XIVb | 6 | 2912727 | 1039841 | 2 |
| Clostridium_difficile_P28_690_innocuum | XVI | 81 | 4323423 | 92019 | 15 |
| Clostridium_innocuum_2789STDY5834853_341 | XVI | 45 | 4283273 | 225726 | 7 |
| Clostridium_innocuum_2959_96 | XVI | 7 | 4803668 | 1547824 | 2 |
| Clostridium_innocuum_NLAE_zl_C381_546 | XVI | 56 | 4232065 | 181667 | 9 |

**Table S2.** **Conserved proteins found among the genomes listed in Table S1**. These genes were used to construct the species tree shown in Figure 1, best amino acid substitution models calculated for each partition in the correspondent matrix are listed.

| **Protein Family** | **Annotation** | **Length** | **Start** | **End** | **Model** |
| --- | --- | --- | --- | --- | --- |
| 1 | Pyruvate:ferredoxin (flavodoxin) oxidoreductase | 1181 | 1 | 1181 | LG+F+I+G4 |
| 2 | Valine--tRNA ligase | 885 | 1182 | 2066 | LG+F+I+G4 |
| 3 | Polyribonucleotide nucleotidyltransferase | 707 | 2067 | 2773 | LG+F+I+G4 |
| 4 | Putative RNA-binding protein | 716 | 2774 | 3489 | LG+F+I+G4 |
| 5 | Aspartate-tRNA ligase | 590 | 3488 | 4079 | LG+I+G4 |
| 6 | Methionine-tRNA ligase | 646 | 4080 | 4725 | LG+F+I+G4 |
| 7 | Excinuclease ABC subunit UvrB | 656 | 4726 | 5381 | LG+F+I+G4 |
| 8 | tRNA uridine-5-carboxymethylaminomethyl(34) synthesis enzyme MnmG | 626 | 5382 | 6007 | LG+F+I+G4 |
| 9 | Elongation factor 4 | 600 | 6008 | 6607 | LG+F+I+G4 |
| 10 | Molecular chaperone DnaK | 616 | 6608 | 7223 | LG+F+I+G4 |
| 11 | Formate-tetrahydrofolate ligase | 557 | 7224 | 7780 | LG+I+G4 |
| 12 | Asparagine--tRNA ligase | 463 | 7781 | 8243 | LG+F+I+G4 |
| 13 | Nicotinate phosphoribosyltransferase | 474 | 8244 | 8717 | LG+F+I+G4 |
| 14 | Chaperonin GroEL | 542 | 8718 | 9259 | LG+F+I+G4 |
| 15 | Ribonuclease Y | 520 | 9260 | 9779 | LG+I+G4 |
| 16 | Methionine adenosyltransferase | 389 | 9780 | 10168 | LG+F+I+G4 |
| 17 | Recombinase RecA | 345 | 10169 | 10513 | LG+F+I+G4 |
| 18 | Flavodoxin-dependent (E)-4-hydroxy-3-methylbut-2-enyl-diphosphate synthase | 350 | 10514 | 10863 | LG+F+I+G4 |
| 19 | 4-hydroxy-tetrahydrodipicolinate synthase | 296 | 10864 | 11159 | LG+I+G4 |
| 20 | Peptide chain release factor 1 | 356 | 11160 | 11515 | LG+I+G4 |
| 21 | Holliday junction branch migration DNA helicase RuvB | 334 | 11516 | 11849 | LG+I+G4 |
| 22 | 6-phosphofructokinase | 319 | 11850 | 12168 | LG+I+G4 |
| 23 | Uracil phosphoribosyltransferase | 209 | 12169 | 12377 | LG+I+G4 |
| 24 | ATP-dependent Clp endopeptidase, proteolytic subunit ClpP | 194 | 12378 | 12571 | LG+I+G4 |
| 25 | 30S ribosomal protein S15 | 86 | 12572 | 12657 | LG+F+I+G4 |
| 26 | 50S ribosomal protein L21 | 103 | 12658 | 12760 | LG+I+G4 |
| 27 | 30S ribosomal protein S16 | 76 | 12761 | 12836 | LG+I+G4 |

**Table S3.** **Conserved proteins found among the 179 genomes with N50>600 kbp listed in Table S1**. These genes were used to construct the species tree shown in Supplementary Figure 16, best amino acid substitution models calculated for each partition in the correspondent matrix are listed.

| **Protein family** | **Annotation** | **Length** | **Start** | **End** | **Model** |
| --- | --- | --- | --- | --- | --- |
| 1 | DNA-directed RNA polymerase beta subunit (EC 2.7.7.6) | 1196 | 1 | 1196 | LG+I+G4 |
| 2 | DNA-directed RNA polymerase beta' subunit (EC 2.7.7.6) | 1134 | 1197 | 2330 | LG+I+G4 |
| 3 | Pyruvate-flavodoxin oxidoreductase (EC 1.2.7.-) | 1188 | 2331 | 3518 | LG+F+I+G4 |
| 4 | Excinuclease ABC subunit A | 533 | 3519 | 4051 | LG+F+I+G4 |
| 5 | Valyl-tRNA synthetase (EC 6.1.1.9) | 523 | 4052 | 4574 | LG+F+I+G4 |
| 6 | Alanyl-tRNA synthetase (EC 6.1.1.7) | 106 | 4575 | 4680 | WAG+I+G4 |
| 7 | ATP-dependent Clp protease, ATP-binding subunit ClpC | 799 | 4681 | 5479 | LG+F+I+G4 |
| 8 | DNA gyrase subunit A (EC 5.99.1.3) | 817 | 5480 | 6296 | LG+F+I+G4 |
| 9 | Transcription accessory protein (S1 RNA-binding domain) | 469 | 6297 | 6765 | LG+F+I+G4 |
| 10 | Translation elongation factor G | 686 | 6766 | 7451 | LG+F+I+G4 |
| 11 | Excinuclease ABC subunit B | 396 | 7452 | 7847 | LG+F+I+G4 |
| 12 | tRNA uridine 5-carboxymethylaminomethyl modification enzyme GidA | 619 | 7848 | 8466 | LG+F+I+G4 |
| 13 | Lysyl-tRNA synthetase (class II) (EC 6.1.1.6) | 497 | 8467 | 8963 | LG+I+G4 |
| 14 | Phosphoglycerate kinase (EC 2.7.2.3) | 390 | 8964 | 9353 | LG+F+I+G4 |
| 15 | Aspartyl-tRNA synthetase (EC 6.1.1.12) | 563 | 9354 | 9916 | LG+I+G4 |
| 16 | Chaperone protein DnaK | 605 | 9917 | 10521 | LG+F+I+G4 |
| 17 | Translation elongation factor LepA | 371 | 10522 | 10892 | LG+I+G4 |
| 18 | Formate--tetrahydrofolate ligase (EC 6.3.4.3) | 473 | 10893 | 11365 | LG+F+I+G4 |
| 19 | Heat shock protein 60 family chaperone GroEL | 526 | 11366 | 11891 | LG+F+I+G4 |
| 20 | GMP synthase [glutamine-hydrolyzing], amidotransferase subunit (EC 6.3.5.2) | 508 | 11892 | 12399 | LG+F+I+G4 |
| 21 | Ribonuclease Y | 478 | 12400 | 12877 | LG+I+G4 |
| 22 | Asparaginyl-tRNA synthetase (EC 6.1.1.22) | 414 | 12878 | 13291 | LG+I+G4 |
| 23 | Glucose-6-phosphate isomerase (EC 5.3.1.9) | 446 | 13292 | 13737 | LG+F+I+G4 |
| 24 | GTP-binding protein EngA | 436 | 13738 | 14173 | LG+I+G4 |
| 25 | Enolase (EC 4.2.1.11) | 413 | 14174 | 14586 | LG+I+G4 |
| 26 | Signal recognition particle, subunit Ffh SRP54 (TC 3.A.5.1.1) | 435 | 14587 | 15021 | LG+F+I+G4 |
| 27 | RNA polymerase sigma factor RpoD | 333 | 15022 | 15354 | LG+I+G4 |
| 28 | ATP-dependent Clp protease ATP-binding subunit ClpX | 380 | 15355 | 15734 | LG+I+G4 |
| 29 | Acetate kinase (EC 2.7.2.1) | 385 | 15735 | 16119 | LG+F+I+G4 |
| 30 | Cysteine desulfurase (EC 2.8.1.7) | 377 | 16120 | 16496 | LG+F+I+G4 |
| 31 | S-adenosylmethionine synthetase (EC 2.5.1.6) | 389 | 16497 | 16885 | LG+I+G4 |
| 32 | RecA protein | 335 | 16886 | 17220 | LG+I+G4 |
| 33 | S-adenosylmethionine:tRNA ribosyltransferase-isomerase | 340 | 17221 | 17560 | LG+F+I+G4 |
| 34 | 1-hydroxy-2-methyl-2-(E)-butenyl 4-diphosphate synthase (EC 1.17.7.1) | 350 | 17561 | 17910 | LG+F+I+G4 |
| 35 | GTP-binding and nucleic acid-binding protein YchF | 365 | 17911 | 18275 | LG+I+G4 |
| 36 | Peptide chain release factor 2 | 308 | 18276 | 18583 | LG+I+G4 |
| 37 | Peptide chain release factor 1 | 354 | 18584 | 18937 | LG+I+G4 |
| 38 | TsaD/Kae1/Qri7 protein, required for threonylcarbamoyladenosine t(6)A37 formation in tRNA | 265 | 18938 | 19202 | LG+I+G4 |
| 39 | Holliday junction DNA helicase RuvB | 328 | 19203 | 19530 | LG+I+G4 |
| 40 | GTP-binding protein Era | 294 | 19531 | 19824 | LG+F+I+G4 |
| 41 | DNA-directed RNA polymerase alpha subunit (EC 2.7.7.6) | 315 | 19825 | 20139 | LG+I+G4 |
| 42 | Ribosomal large subunit pseudouridine synthase D (EC 4.2.1.70) | 301 | 20140 | 20440 | cpREV+I+G4 |
| 43 | rRNA small subunit methyltransferase H | 311 | 20441 | 20751 | LG+F+I+G4 |
| 44 | Signal peptidase-like protein | 264 | 20752 | 21015 | LG+I+G4 |
| 45 | Predicted P-loop-containing kinase | 283 | 21016 | 21298 | LG+I+G4 |
| 46 | LSU ribosomal protein L2p (L8e) | 261 | 21299 | 21559 | LG+I+G4 |
| 47 | Transmembrane component of general energizing module of ECF transporters | 259 | 21560 | 21818 | LG+F+I+G4 |
| 48 | SSU ribosomal protein S3p (S3e) | 213 | 21819 | 22031 | LG+I+G4 |
| 49 | SSU ribosomal protein S2p (SAe) | 202 | 22032 | 22233 | LG+I+G4 |
| 50 | Septum site-determining protein MinD | 244 | 22234 | 22477 | LG+I+G4 |
| 51 | RNA polymerase sporulation specific sigma factor SigE | 221 | 22478 | 22698 | LG+I+G4 |
| 52 | RNA polymerase sporulation specific sigma factor SigG | 256 | 22699 | 22954 | LG+G4 |
| 53 | LSU ribosomal protein L1p (L10Ae) | 229 | 22955 | 23183 | LG+I+G4 |
| 54 | Redox-sensitive transcriptional regulator (AT-rich DNA-binding protein) | 167 | 23184 | 23350 | LG+I+G4 |
| 55 | LSU ribosomal protein L3p (L3e) | 207 | 23351 | 23557 | LG+I+G4 |
| 56 | Uracil phosphoribosyltransferase (EC 2.4.2.9) | 209 | 23558 | 23766 | LG+G4 |
| 57 | LSU ribosomal protein L4p (L1e) | 169 | 23767 | 23935 | LG+I+G4 |
| 58 | SSU ribosomal protein S4p (S9e) | 101 | 23936 | 24036 | LG+I+G4 |
| 59 | ATP-dependent Clp protease proteolytic subunit (EC 3.4.21.92) | 192 | 24037 | 24228 | LG+I+G4 |
| 60 | Translation initiation factor 3 | 123 | 24229 | 24351 | LG+I+G4 |
| 61 | Translation elongation factor P | 185 | 24352 | 24536 | LG+G4 |
| 62 | Transcription antitermination protein NusG | 171 | 24537 | 24707 | LG+I+G4 |
| 63 | LSU ribosomal protein L6p (L9e) | 179 | 24708 | 24886 | LG+I+G4 |
| 64 | LSU ribosomal protein L5p (L11e) | 179 | 24887 | 25065 | LG+I+G4 |
| 65 | SSU ribosomal protein S7p (S5e) | 177 | 25066 | 25242 | LG+I+G4 |
| 66 | SSU ribosomal protein S5p (S2e) | 73 | 25243 | 25315 | LG+G4 |
| 67 | tmRNA-binding protein SmpB | 151 | 25316 | 25466 | LG+I+G4 |
| 68 | Transcription elongation factor GreA | 158 | 25467 | 25624 | LG+I+G4 |
| 69 | LSU ribosomal protein L15p (L27Ae) | 92 | 25625 | 25716 | LG+I+G4 |
| 70 | SSU ribosomal protein S8p (S15Ae) | 132 | 25717 | 25848 | LG+I+G4 |
| 71 | LSU ribosomal protein L14p (L23e) | 122 | 25849 | 25970 | LG+I+G4 |
| 72 | LSU ribosomal protein L20p | 117 | 25971 | 26087 | LG+G4 |
| 73 | LSU ribosomal protein L21p | 101 | 26088 | 26188 | LG+I+G4 |
| 74 | SSU ribosomal protein S15p (S13e) | 80 | 26189 | 26268 | LG+G4 |
| 75 | SSU ribosomal protein S17p (S11e) | 67 | 26269 | 26335 | FLU+I+G4 |
| 76 | SSU ribosomal protein S16p | 76 | 26336 | 26411 | LG+G4 |
| 77 | KH domain RNA binding protein YlqC | 50 | 26412 | 26461 | rtREV+G4 |
| 78 | Translation initiation factor 1 | 72 | 26462 | 26533 | LG+G4 |
| 79 | LSU ribosomal protein L11p (L12e) | 90 | 26534 | 26623 | LG+I+G4 |

**Table S4. Distribution of toxin homologues in *Clostridium* species. “**X” indicates that the species has the homolog of the toxin.

|  |  | **Toxins** | | | |
| --- | --- | --- | --- | --- | --- |
|  |  | *C. difficile*  A/B toxins* | *C. perfringens* alpha toxin | *C. septicum* alpha toxin | *C. botulinum/tetani*  toxin |
| Taxonomic group | *C. difficile* | X |  |  |  |
|  | *C. acetobutylicum* | X |  |  |  |
|  | *C. sordelli* | X | X |  |  |
|  | *C. novyi* | X | X | X |  |
|  | *C. haemolyticum* |  | X | X |  |
|  | *C. botulinum C & D* |  | X | X | X |
|  | *C. septicum* |  |  | X |  |
|  | *C. perfringens* |  | X |  |  |
|  | *C. cavendishii* |  | X |  |  |
|  | *C. dakarense* |  | X |  |  |
|  | *C. argentinense* |  | X |  | X |
|  | *C. baratii* |  | X |  | X |
|  | *C. tetani* |  |  |  | X |
|  | *C. butyricum* |  |  |  | X |
|  | *C. botulinum A, B, E & F* |  |  |  | X |

*****TpeL from C. perfringens has been defined homolog of the C. difficile toxins A and B toxins (see Amimoto et al, Microbiology. 2007 153:1198-206) however we excluded them among the C. difficile toxins given the large divergence between them.

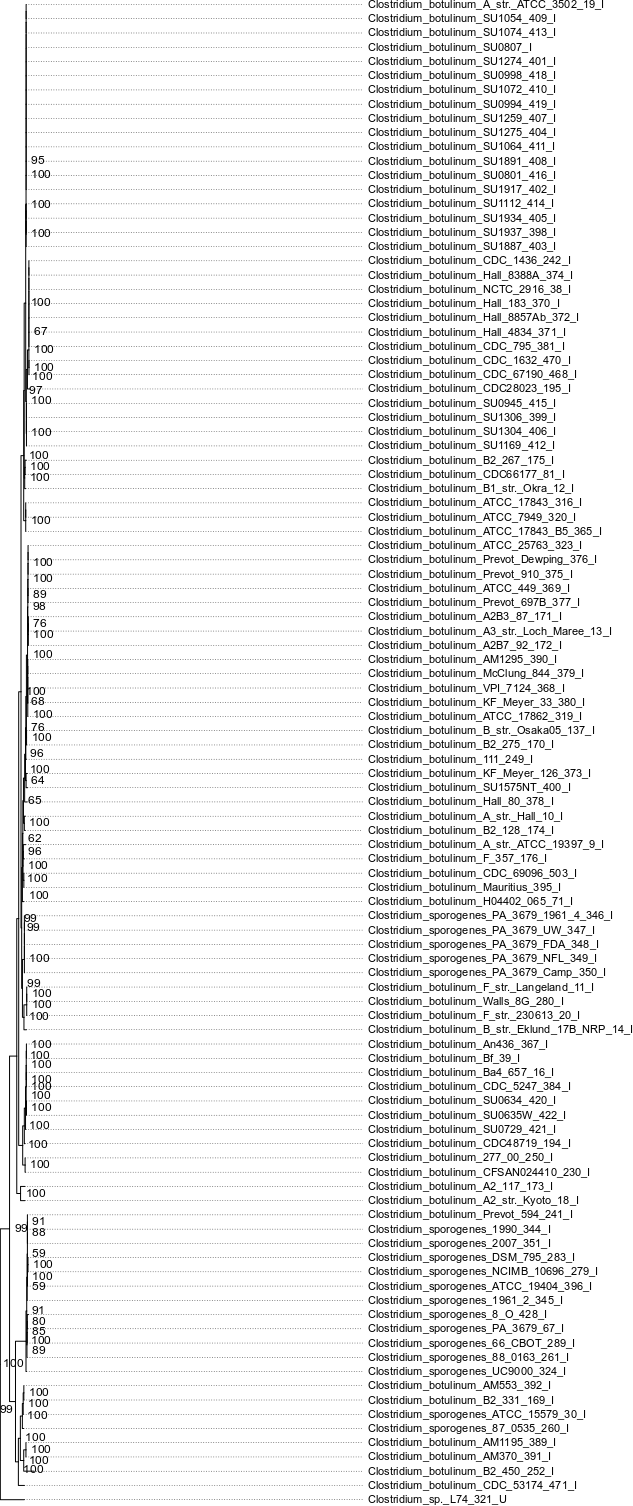
**Figure S1. Zoom-in of subgroup 1 in the species tree shown in Figure 1**

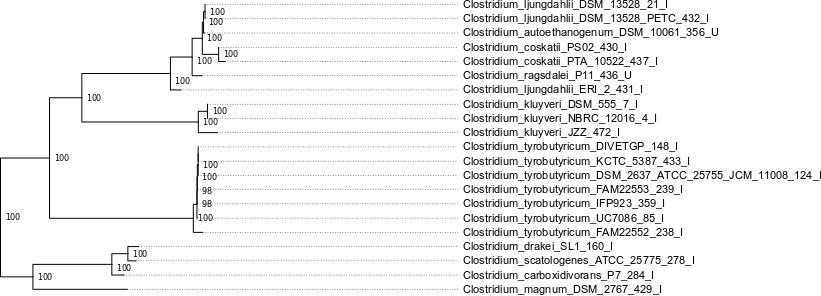

**Figure S2. Zoom-in of subgroup 2 in the species tree shown in Figure 1**

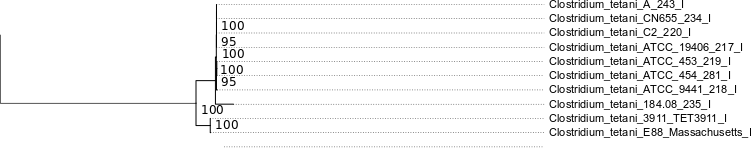

**Figure S3. Zoom-in of subgroup 3 in the species tree shown in Figure 1**

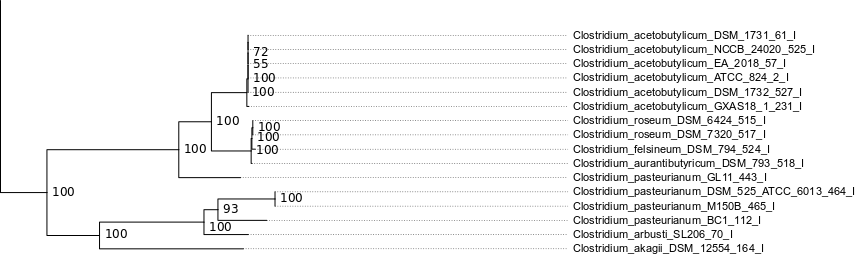

**Figure S4. Zoom-in of subgroup 4 in the species tree shown in Figure 1**

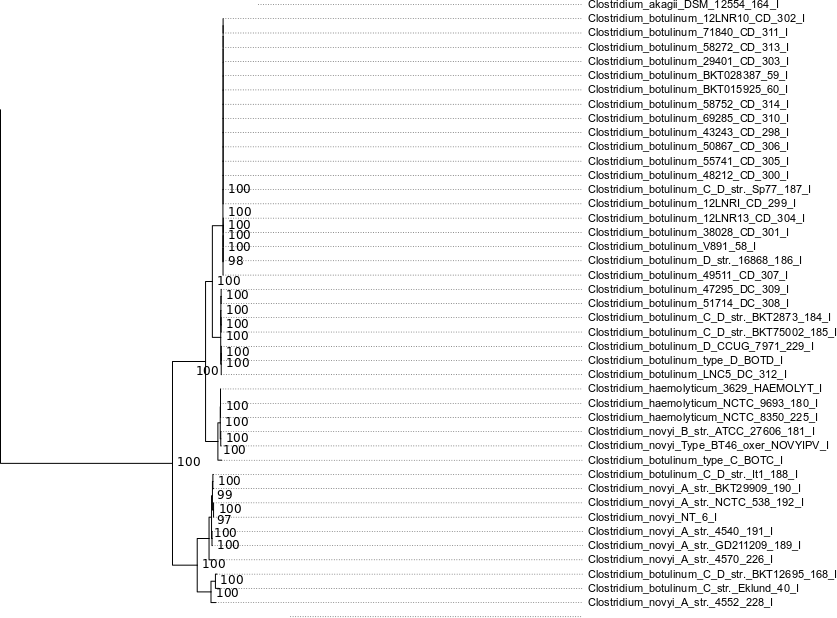

**Figure S5. Zoom-in of subgroup 5 in the species tree shown in Figure 1**

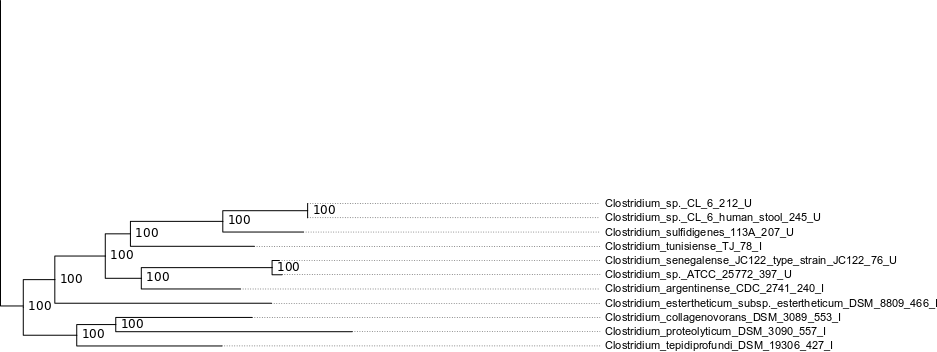

**Figure S6. Zoom-in of subgroup 6 in the species tree shown in Figure 1**

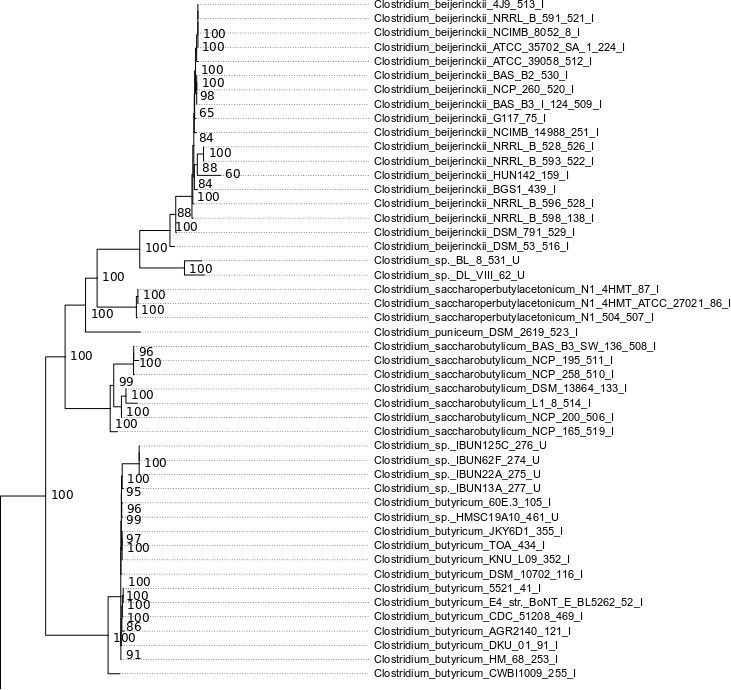
**Figure S7. Zoom-in of subgroup 7 in the species tree shown in Figure 1**

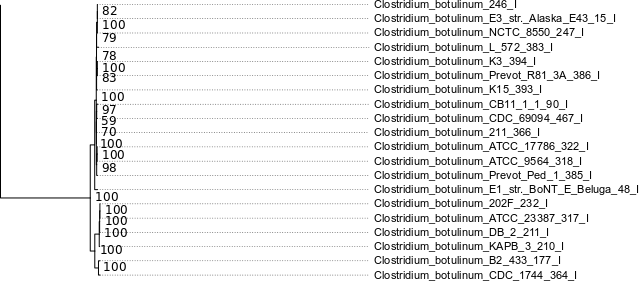

**Figure S8. Zoom-in of subgroup 8 in the species tree shown in Figure 1**

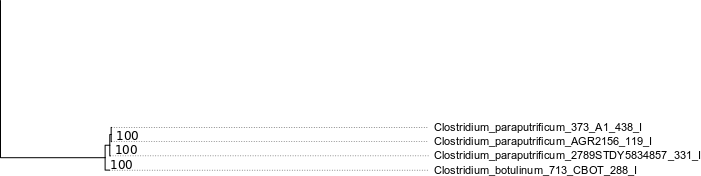

**Figure S9. Zoom-in of subgroup 9 in the species tree shown in Figure 1**

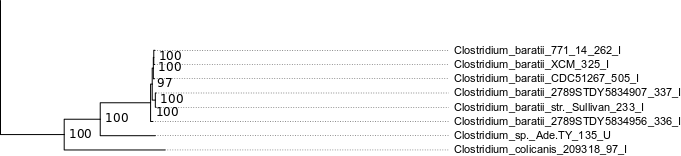

**Figure S10. Zoom-in of subgroup 10 in the species tree shown in Figure 1**

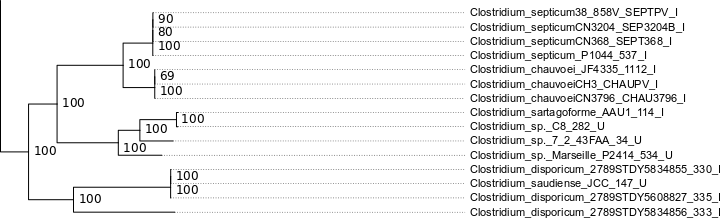

**Figure S11. Zoom-in of subgroup 11 in the species tree shown in Figure 1**

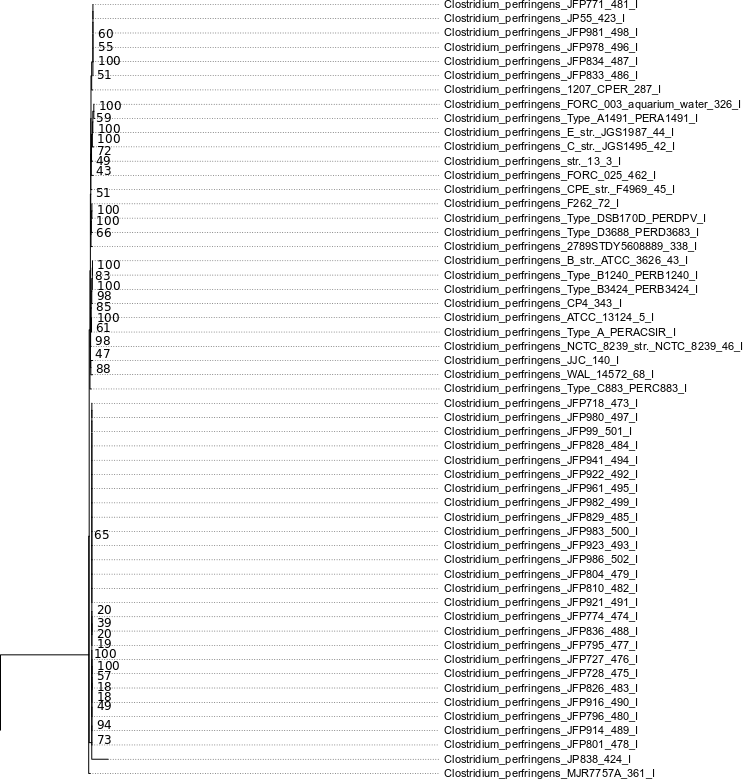

**Figure S12. Zoom-in of subgroup 12 in the species tree shown in Figure 1**

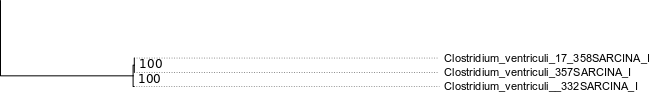

**Figure S13. Zoom-in of subgroup 13 in the species tree shown in Figure 1**

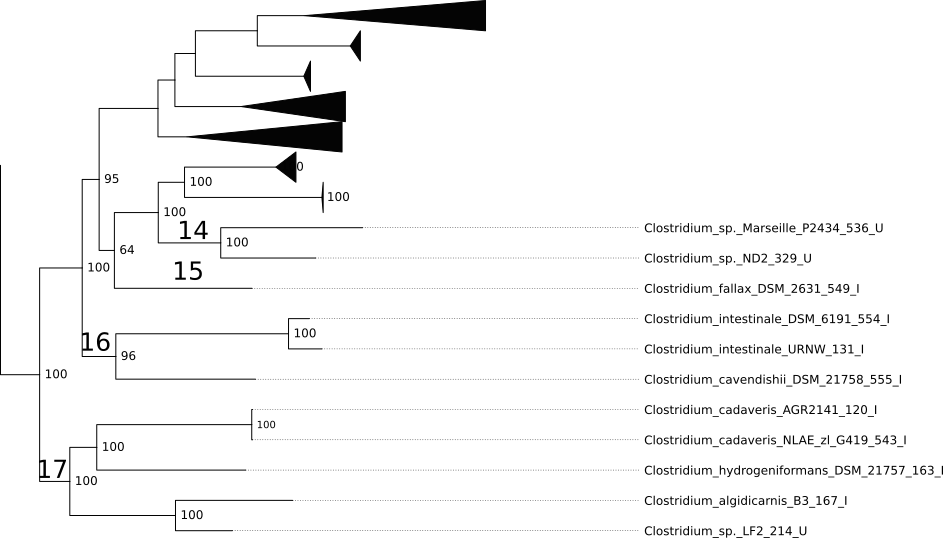
**Figure S14. Zoom-in of subgroups 14-17 in the species tree shown in Figure 1**
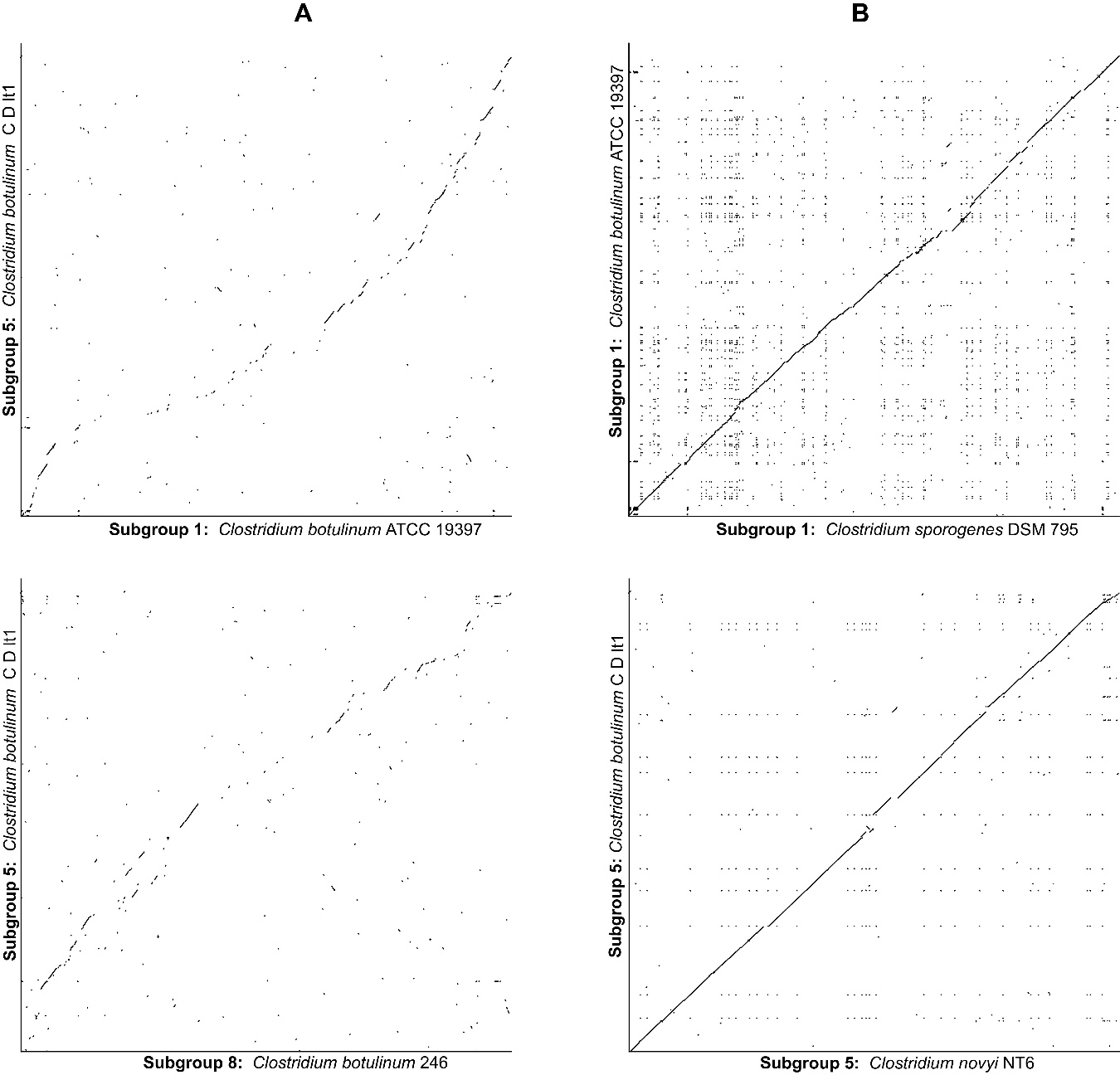

**Figure S15. (A)** Synteny among *C. botulinum* strains from different subgroups and **(B)** other closely related strains under distinct species names. DNA sequences of single contig genomes (except for *C. botulinum* C D lt1) were compared using R2CAT (<https://bibiserv2.cebitec.uni-bielefeld.de/cgcat>). Lines represent homologous regions. Syntenic regions are aligned along the diagonal center between the compared genomes. From this representation is clear that *C. botulinum* from Subgroup 1 and *C. sporogenes*; and *C. botulinum* from subgroup 5 and *C. novyi* (defined as a different species) are more syntenic than *C. botulinum* from subgroups 1, 5 and 8 (defined as the same species).

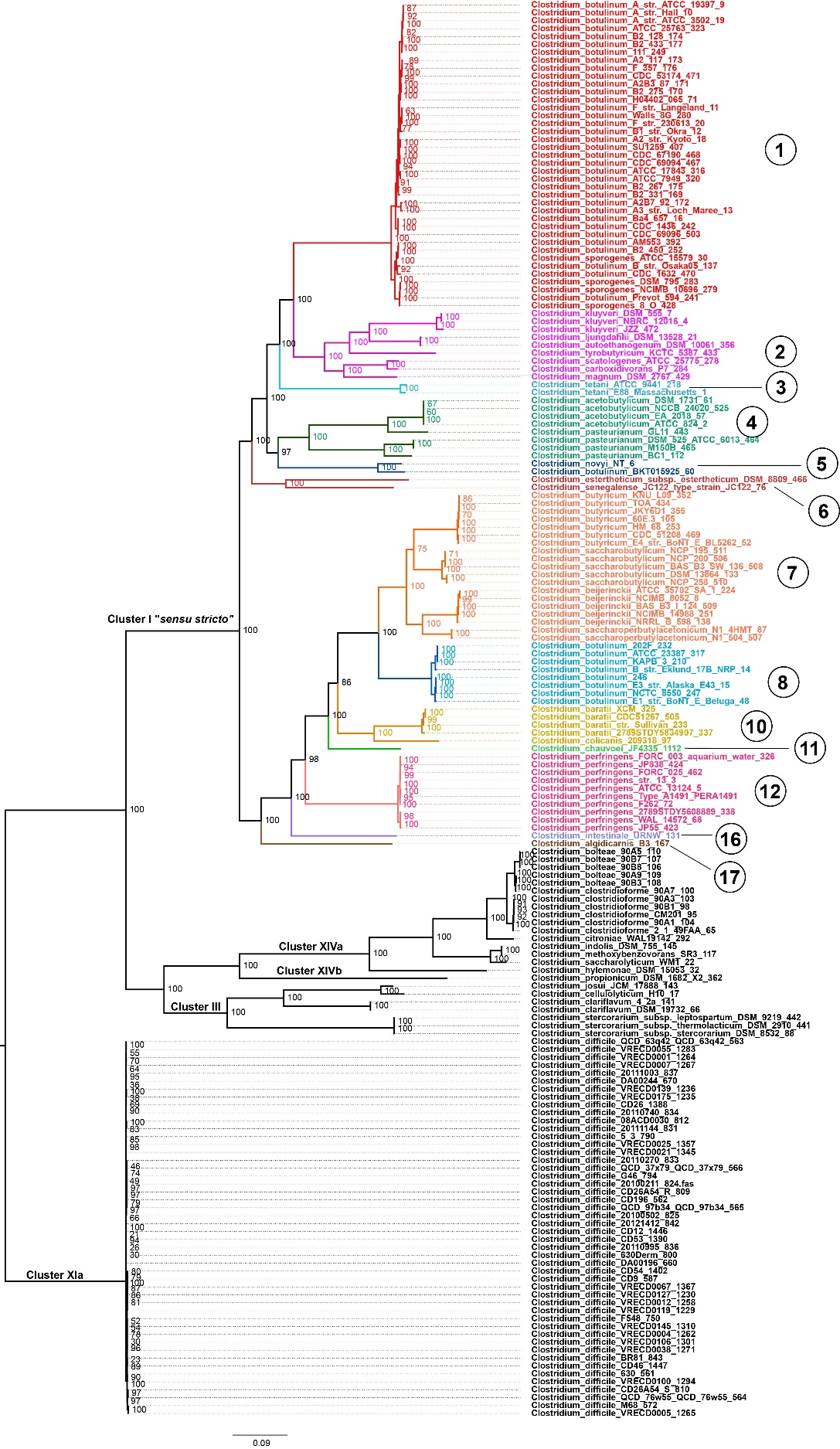

**Figure S16. Second phylogenetic reconstruction of *Clostridium* species.** This phylogeny was constructed using 79 markers conserved across 179 genomes (N50>600,000 and removing unclassified genomes (*Clostridium_sp*) from Supplementary Table S1) deposited in the GenBank database and taxonomically defined as *Clostridium*. Each partition (79 proteins) was aligned independently and manually curated. The best independent evolutionary model was determined, and 10,000 bootstrap replicates were performed. The main clades outside and within the *sensu stricto* group (real clostridia) have been defined as taxonomic subgroups (Table 1 in Manuscript). Branch support is shown at each node. Zoom-in of subgroups of Cluster I are shown in Supplementary Figures S17-S29.

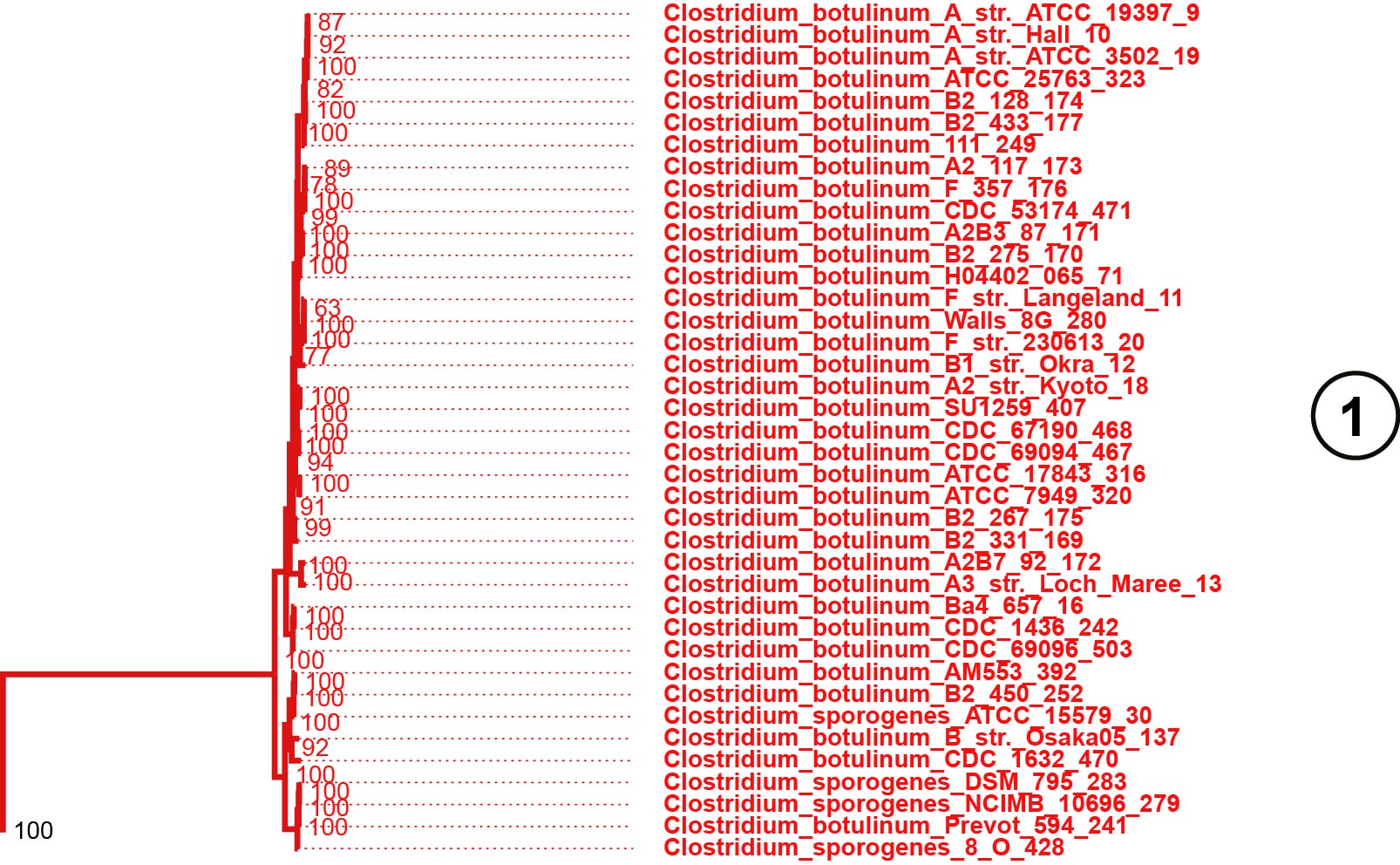

**Figure S17. Zoom-in of subgroup 1 in the species tree shown in Figure S16**

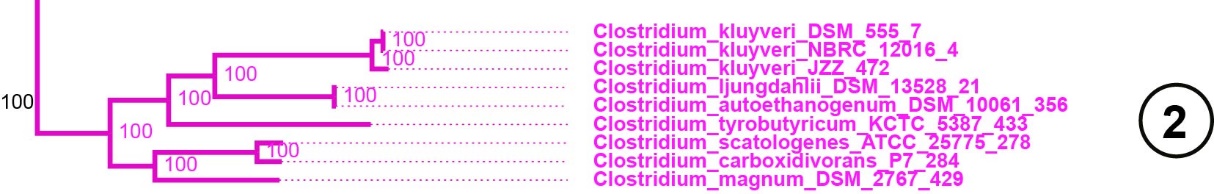

**Figure S18. Zoom-in of subgroup 2 in the species tree shown in Figure S16**

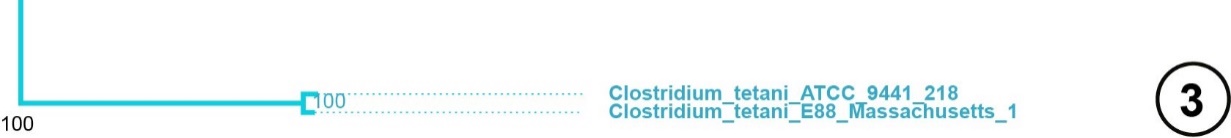

**Figure S19. Zoom-in of subgroup 3 in the species tree shown in Figure S16**

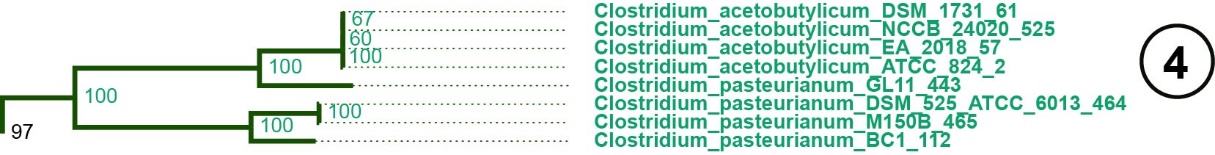

**Figure S20. Zoom-in of subgroup 4 in the species tree shown in Figure S16**

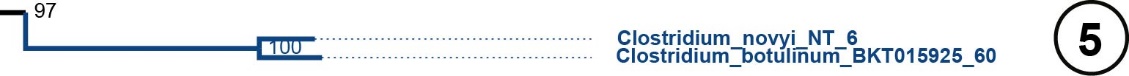

**Figure S21. Zoom-in of subgroup 5 in the species tree shown in Figure S16**

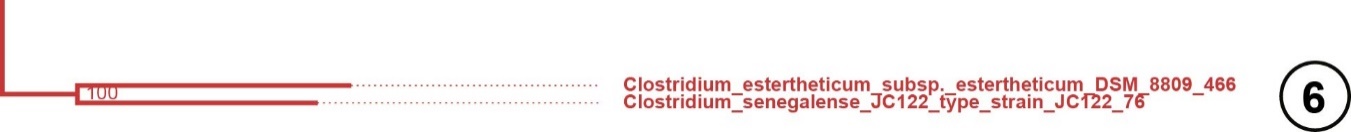

**Figure S22. Zoom-in of subgroup 6 in the species tree shown in Figure S16**

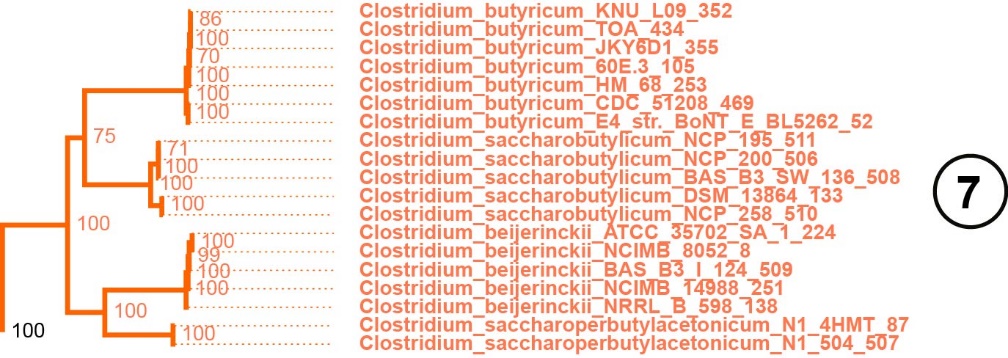

**Figure S23. Zoom-in of subgroup 7 in the species tree shown in Figure S16**

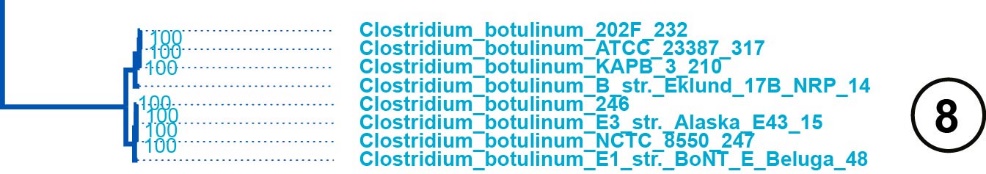

**Figure S24. Zoom-in of subgroup 8 in the species tree shown in Figure S16**

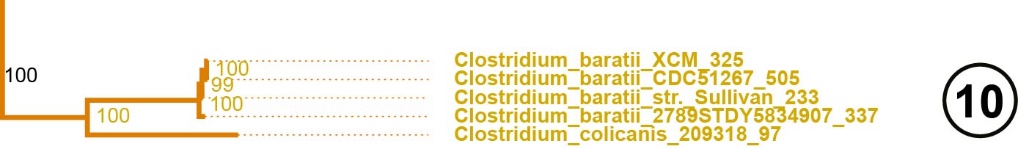

**Figure S25. Zoom-in of subgroup 10 in the species tree shown in Figure S16**

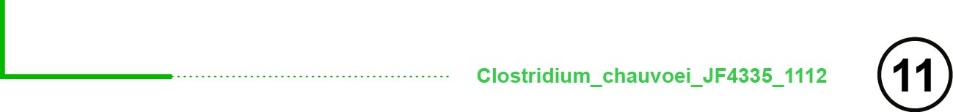

**Figure S26. Zoom-in of subgroup 11 in the species tree shown in Figure S16**

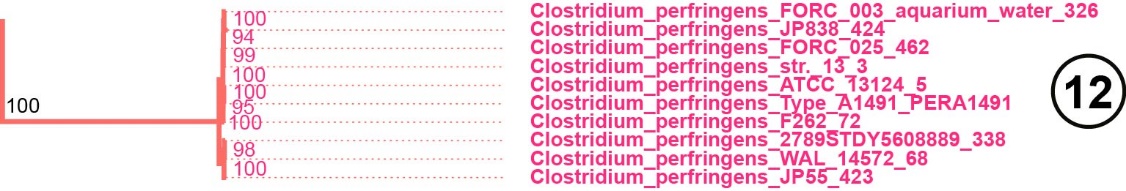

**Figure S27. Zoom-in of subgroup 12 in the species tree shown in Figure S16**

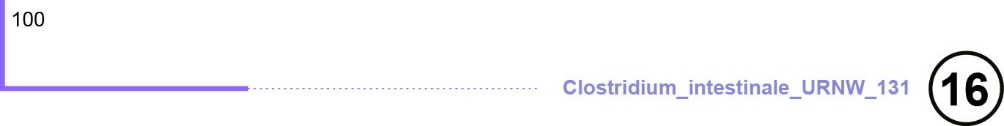

**Figure S28. Zoom-in of subgroup 16 in the species tree shown in Figure S16**

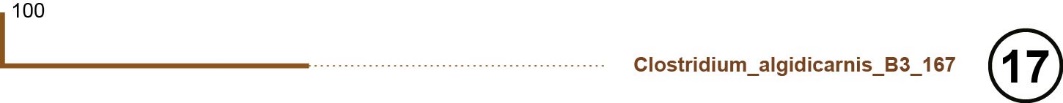

**Figure S29. Zoom-in of subgroup 17 in the species tree shown in Figure S16**

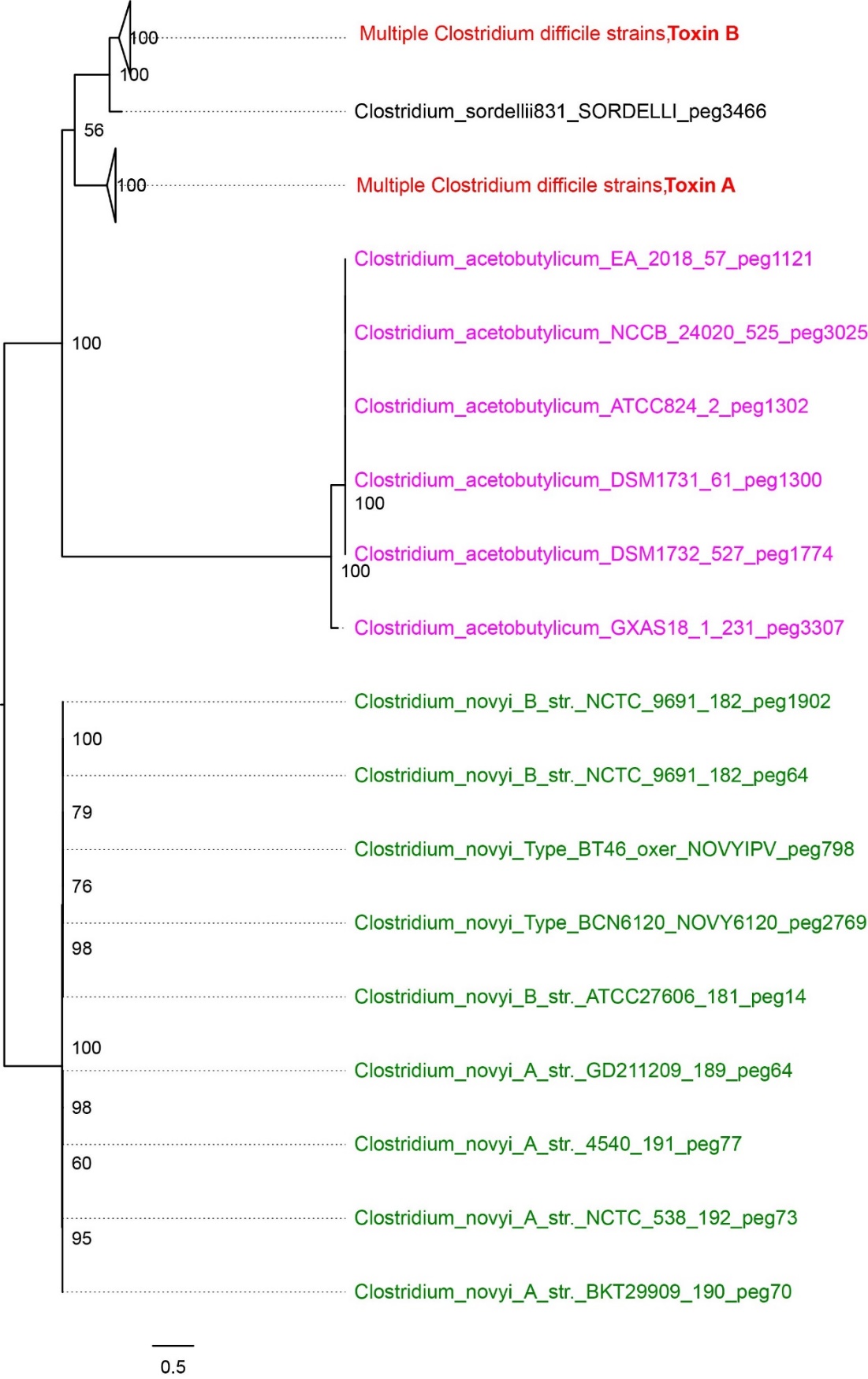

**Figure S30. Phylogenetic reconstruction of *C. difficile* toxins A and B (shown in red) homologous proteins.** Three non-*difficile* strains are included: *C. sordellii*, *C. acetobutylicum* and *C. novyi*.

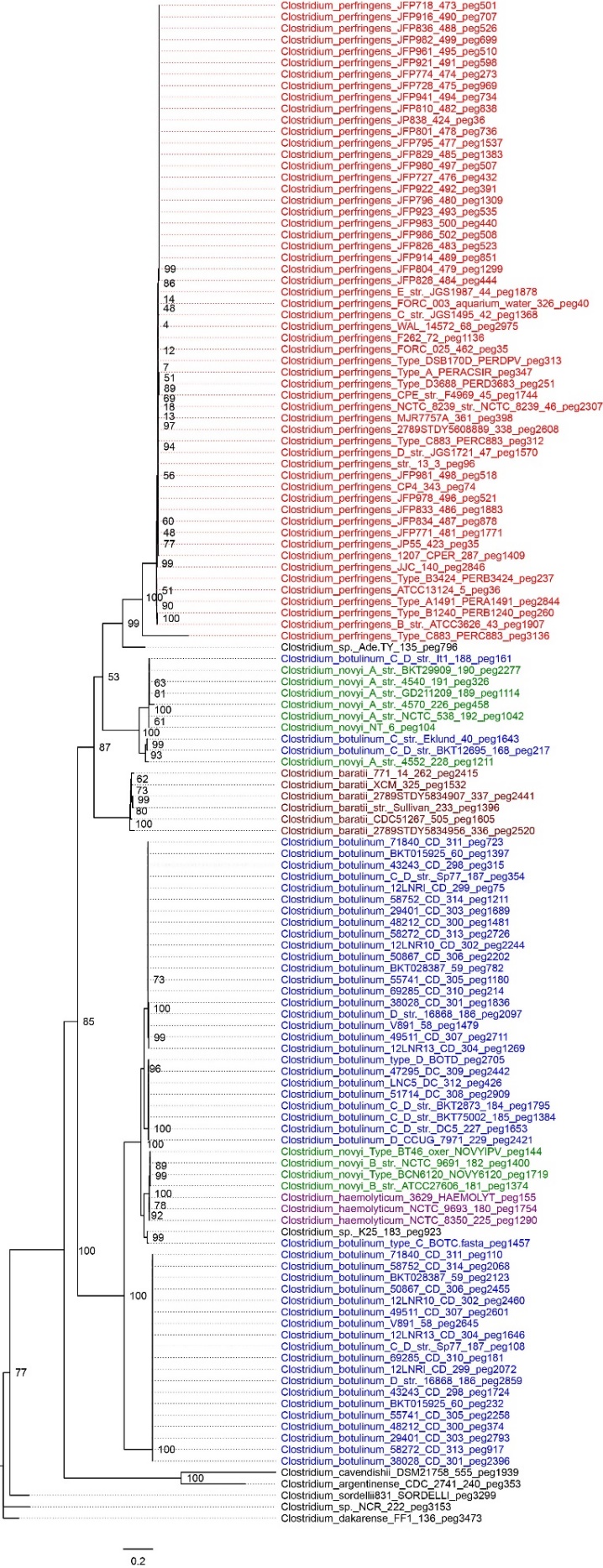

**Figure S31. Phylogenetic reconstruction of *C. perfringens* alpha** **toxin (shown in red) homologous proteins.** Nine non-*perfringens* strains are included: *C. novyi*, *C. botulinum* C and D, *C. baratii*, *C. hemolyticum*, *C. cavendishii*, *C. argentinense*, *C. sordellii* and *C. dakarense*.

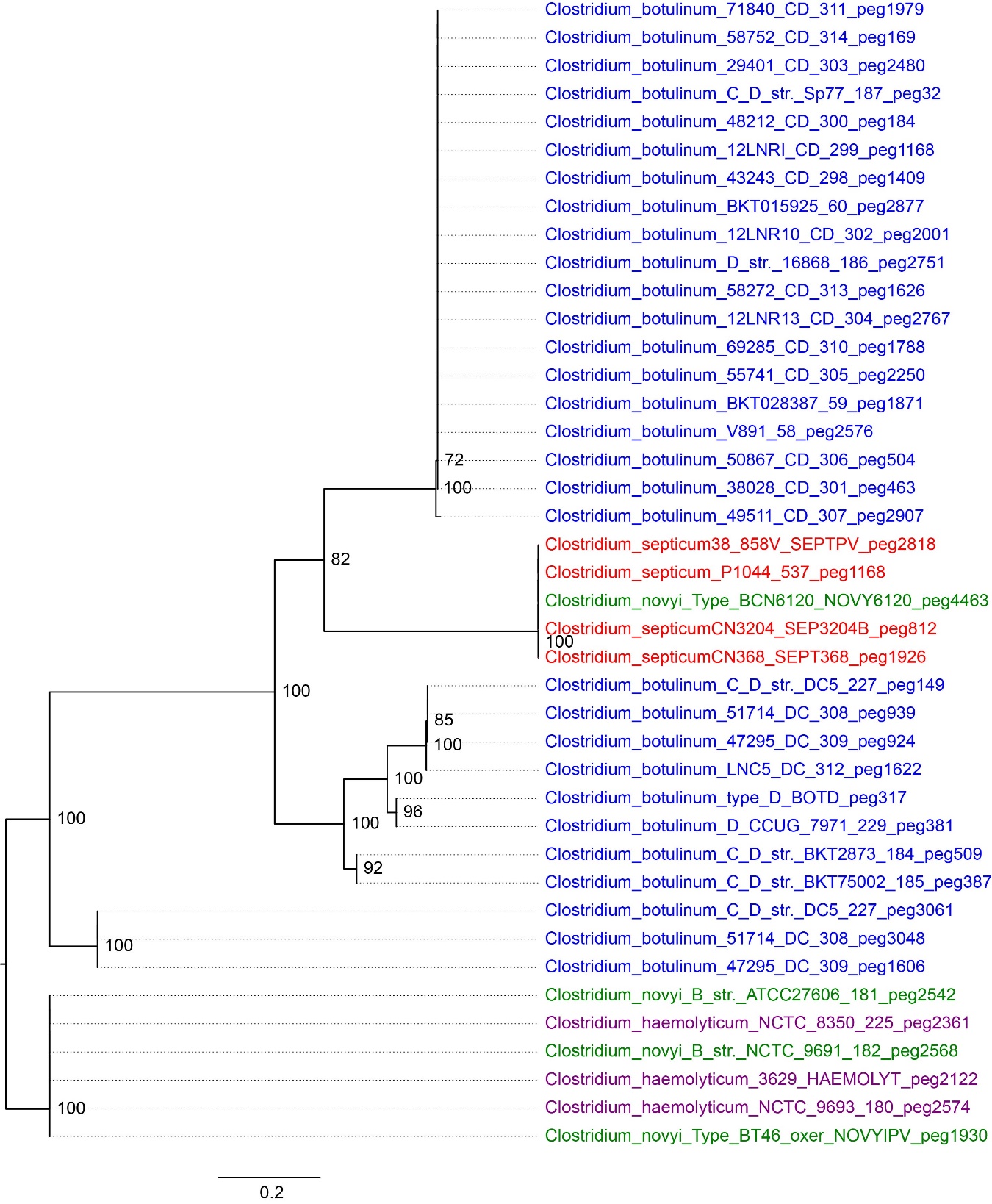

**Figure S32. Phylogenetic reconstruction of *C. septicum* alpha** **toxin (shown in red) homologous proteins.** Four non- *septicum* strains are included: *C. novyi*, *C. haemolyticum* and *C. botulinum* C and D species.

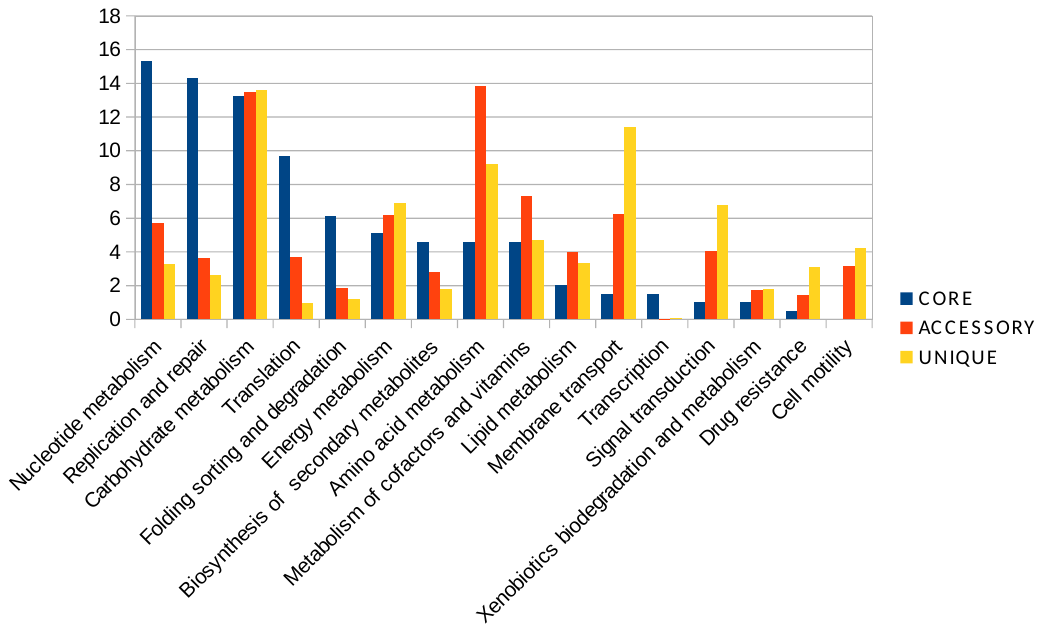

**Figure S33**. **PanGenome analysis of Cluster I strains.** This figure represents the functional content shown as KEGG orthology categories of the core, accessory ad unique genes found in cluster I. Values in the y axis are the percentage of annotated core, accessory or unique functions in each category in the x axis.
